# Probabilistic mouse-human brain map translation through multimodal optimal transport

**DOI:** 10.64898/2026.08.24.746652

**Authors:** Stefan P. Koch, Mario Perales, Tanmoy Sil, Shawn Hiew, Johannes Hartig, Benedikt Weigl, Muthuraman Muthuraman, Florian Lange, Ningfei Li, Andrea A. Kühn, Katharina Breininger, Philip Tovote, Chi Wang Ip, Jens Volkmann, Martin Reich, Philipp Boehm-Sturm, Robert Peach

## Abstract

Cross-species differences in brain organisation complicate the interpretation of mouse experiments and the selection of mouse regions from human imaging findings. Here we present OTTER, which combines functional and structural connectivity, spatial position and comparative anatomy through optimal transport to estimate probabilistic mouse–human brain correspondence. The mapping recovers established regional relationships and supports bidirectional translation of experimental, functional and molecular maps. Assessment of the connectomes used to fit OTTER shows lower mouse-based reconstruction accuracy for human functional connectivity in the expanded association cortex compared to sensory and motor regions. Cross-sectional human Parkinson disease maps yield translated cortical thinning scores that increase across stages in the mouse visceral, gustatory and agranular-insular regions relative to the motor cortex. The corresponding anatomical contrast increases over time in a separate longitudinal MRI analysis following α-synuclein fibril injection in mice. OTTER predicts human maps from mouse experiments and candidate mouse targets from human imaging.

**Editorial summary:** OTTER combines brain connectivity, spatial position and comparative anatomy through optimal transport to estimate probabilistic correspondence between mouse and human brain regions. The mapping generates predicted human maps from mouse experimental responses and identifies candidate mouse regions for testing hypotheses derived from human imaging.

## Introduction

Mouse studies allow targeted recording and manipulation of neural circuits using genetic, viral and optogenetic tools^1^. Relating these experiments to human brain function and disease requires identifying corresponding regions, both to interpret mouse findings and to select mouse targets for hypotheses derived from human imaging. However, mouse and human brains differ in regional size, organisation and connectivity, particularly in the expanded human association cortex^2^. For example, the rodent medial frontal cortex is often used to investigate the human prefrontal function^3,4^, yet rat medial frontal functional-connectivity patterns resemble those of marmoset premotor cortex more closely than those of marmoset dorsolateral prefrontal cortex^5^. Regional correspondence therefore needs to be evaluated in relation to the anatomical and connectional features relevant to each experimental question.

In practice, cross-species correspondences are commonly drawn from comparative anatomical literature and brain atlases. Regions are matched using anatomical position, cytoarchitecture, connectivity and nomenclature^4,8^. These assignments provide a foundation for comparison, but do not by themselves specify how a brain map should be translated between species. For example, the mouse caudoputamen corresponds to both the human caudate and putamen^9^. This regional relationship alone leaves open how measurements at different locations within the caudoputamen should contribute to a translated human map. Quantitative mappings can specify weights between locations for translating brain maps and assessing connectivity correspondence.

Computational approaches estimate correspondence using connectivity and molecular information. Connectivity-based methods compare regions through their relationships with the rest of the brain^10^. Applications include comparisons between the human and macaque cortex^11–13^ and between rodents and primates^5^, using connectivity blueprints and joint embeddings^11,14^. These approaches also support transfer of cortical maps between species^11,14^. Molecular approaches use cross-species similarities in gene expression^9,15,16^. TransBrain combines transcriptomic embeddings with species-specific structural connectivity and anatomical hierarchy to translate brain-wide phenotypes between mouse and human^16^, while recent work also uses gene-expression representations for voxel-wise translation of disease maps^102^. We therefore asked whether a whole-brain mapping fitted to functional and structural connectivity, with spatial and anatomical guidance, could better relate mouse experimental maps to human brain organisation and identify candidate mouse regions from human imaging.

Here we present OTTER (Optimal Transport for Translation of Evolutionary Relatives), a method that uses multimodal optimal transport^19,20,21,22^ to estimate correspondence between mouse and human brain parcels. The resulting coupling is a matrix of correspondence weights between pairs of sampled brain locations. OTTER fits these weights using functional and structural connectivity, spatial position, anatomical landmarks selected from a comparative atlas^23^ and curated regional correspondences (Fig. 1b–d). Normalising each mouse parcel’s weights to sum to one gives a distribution over human parcels, allowing correspondence to extend across several regions (Fig. 1a). This coupling translates brain maps in either direction through weighted averaging and supports assessments of regional correspondence and connectivity reconstruction.

**Figure 1.**
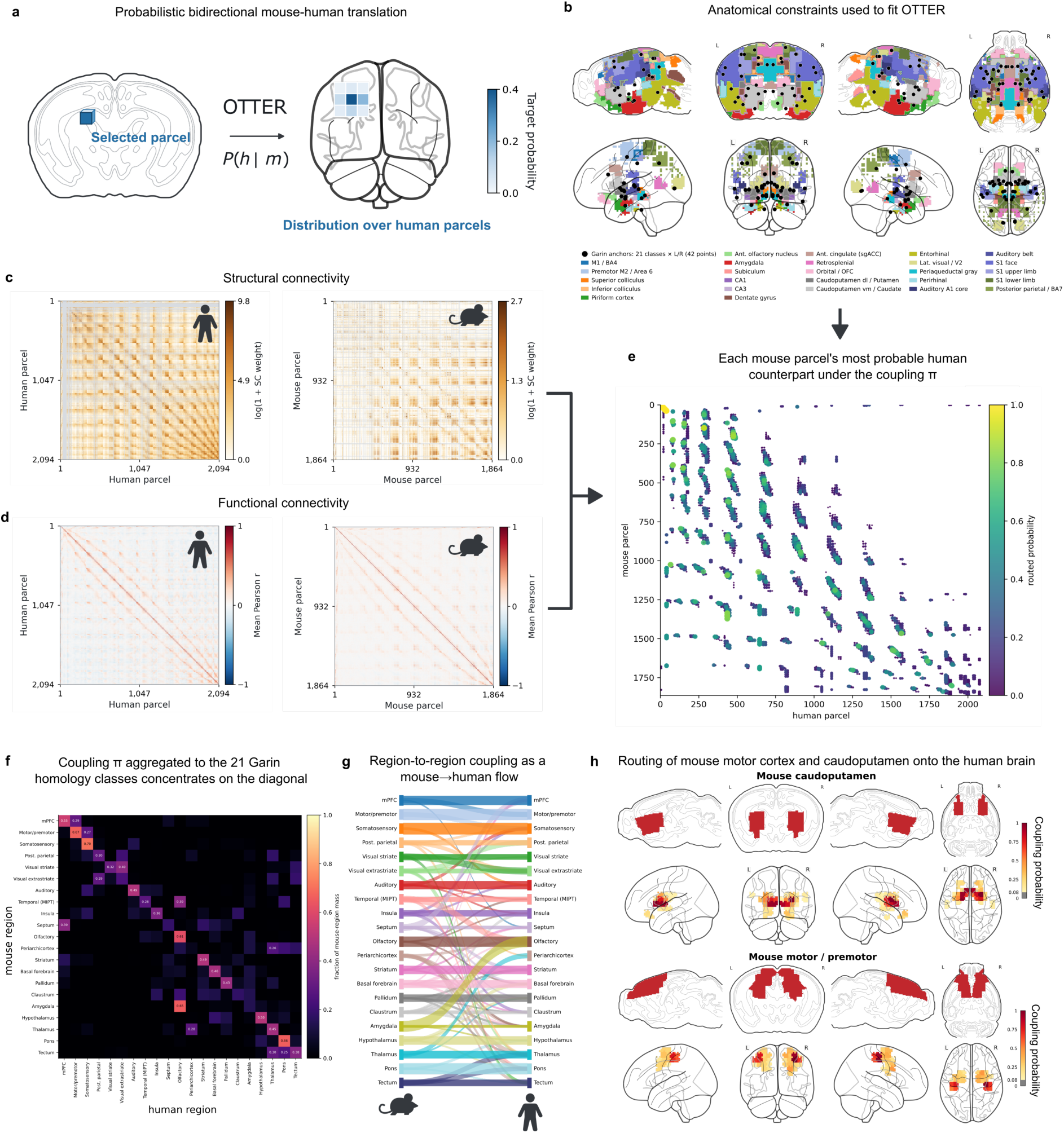
OTTER estimates probabilistic mouse–human correspondence from anatomy and connectivity. **a**, Schematic of a selected mouse parcel’s probability distribution over human parcels, obtained by normalising its coupling weights to sum to one. Parcel geometry and probabilities are illustrative. **b**, Anatomical constraints used in fitting (mouse, top; human, bottom): 42 point anchors per species (black dots), representing 21 bilateral anatomical classes selected from the atlas of Garin et al. ^23^, and 26 curated regional correspondences (coloured regions; Supplementary Table 1). Point-anchor pairs also define the coordinate transformation used to calculate the spatial cost. **c**, Structural-connectivity matrices for human (left; 2,094 parcels) and mouse (right; 1,864 parcels), shown as log(1 + connection weight). Human weights are streamline counts and mouse weights are normalised projection volumes, with separate colour scales. Grey marks unavailable structural-connectivity coverage. **d**, Group-mean functional-connectivity matrices using the same parcel order as c within each species, showing mean Pearson correlations from 113 human participants (left) and 105 mice (right). **e**, Highest-weight human target for each mouse parcel. Each point shows one mouse parcel and its selected human target; colour indicates the target’s probability within the mouse parcel’s distribution (median 0.31; above 0.5 for 20% of mouse parcels). **f**, Coupling aggregated across 21 anatomical classes, assigning each parcel to its nearest point anchor in the same hemisphere and normalising each mouse-class row to sum to one. **g**, The same class-level fractions shown as ribbons from mouse (left) to human (right); wider ribbons indicate larger fractions, with fractions above 0.06 displayed. **h**, Selected mouse parcel sets (red, upper row of each example) and their human correspondence-weight maps (lower row), obtained by summing weights across source mouse parcels and scaling each map to its own maximum.

We evaluate OTTER against established regional correspondences and test bidirectional translation of experimental, functional and molecular maps, including comparisons with TransBrain. Translating the activation map evoked by optogenetic stimulation of mouse agranular insula produced a human prediction enriched in salience/ventral-attention territories. An internal assessment of the connectomes used to fit OTTER shows less accurate mouse-based reconstruction of human functional connectivity in the expanded association cortex than in sensory and motor territories. Translations of human Alzheimer disease phenotype maps support distinct prespecified mouse target sets, and symptom-specific stimulation target maps for depression prioritise different candidate regions. In Parkinson disease, translated cross-sectional maps of human cortical thinning show an overall stage-related increase in scores for visceral, gustatory and agranular-insular regions relative to the motor cortex. This pattern is also present in an external human cohort; the corresponding anatomical contrast increases over time in a separate longitudinal α-synuclein mouse MRI analysis. These results provide an anatomical basis for relating mouse experimental maps to human brain organisation and selecting mouse brain regions to test hypotheses derived from human imaging.

## Results

### OTTER learns a probabilistic mouse–human coupling

To estimate correspondence between mouse and human brain parcels, we fitted a coupling matrix, π, using fused Gromov–Wasserstein optimal transport. The optimisation combined within-species functional and structural connectivity relationships with cross-species spatial costs and soft anatomical constraints to assign correspondence weights to mouse–human parcel pairs (Fig. 1a–d; Methods). Anatomical guidance comprised 42 point anchors per species, representing 21 bilateral anatomical classes selected from the atlas of Garin et al.^23^, and 26 regional correspondences curated from comparative neuroanatomical studies (Supplementary Table 1). The point-anchor pairs also defined a smooth transformation from mouse to human coordinates for calculating the spatial cost. We used a semirelaxed formulation^24^ that fixed the same total weight for each mouse parcel while leaving the weight received by each human parcel unconstrained. We also examined a refit with fixed human parcel totals and unconstrained mouse totals (Extended Data Fig. 9).

The fitted coupling linked 1,864 mouse parcels to 2,094 human parcels. Normalising each mouse parcel’s correspondence weights to sum to one gave a distribution over human parcels. These distributions generally spread weight across several targets: the highest-weighted parcel had a median weight of 0.31 and received more than half of the total weight for 20% of mouse parcels (Fig. 1e). To summarise correspondence at the regional level, we grouped parcels by their nearest point anchor in the same hemisphere and aggregated weights across the 21 paired classes. The mean fraction assigned to the corresponding human class was 0.40, approximately 8.5 times the mean of 0.0476 under uniform parcel-to-parcel weights (Fig. 1f,g). For region-level queries, we summed correspondence weights over mouse parcels within a selected region and ranked human parcels by their received mass. Queries for the mouse motor/premotor and striatal classes, for example, assigned the highest weights to the human motor cortex and striatum, respectively (Fig. 1h).

To assess sensitivity to cohort sampling, we refitted OTTER using functional connectivity estimated from random halves of the human and mouse cohorts. Regional correspondence was evaluated using 19 mouse–human region pairs adapted from Beauchamp et al.^9^, quantifying how well coupling mass received from the corresponding mouse region distinguished parcels in the expected human region from other human parcels (area under the receiver operating characteristic curve, AUROC; Methods). The two refits gave AUROCs of 0.896 and 0.903, compared with 0.899 for the full cohorts; their coupling weights were correlated across corresponding matrix entries (r = 0.995), and 81% of mouse parcels retained the same highest-weight human target across the two halves. Removing structural connectivity (AUROC 0.884) or functional connectivity (AUROC 0.885) changed 37% and 59% of the highest-weight targets, respectively, relative to the combined model. Regional scores therefore changed minimally, whereas removing a connectivity modality altered individual parcel assignments more than the cohort split did (Extended Data Fig. 2a–c).

We also assessed the spatial organisation of the fitted coupling by comparing pairwise distances between mouse parcels with distances between their correspondence-weighted human centroids. These distances were correlated (r = 0.53), compared with correlations near zero after permuting the coupling rows.

### Connectivity, spatial structure and anatomical supervision make complementary contributions

To assess how the cost components affected regional correspondence, we refitted OTTER while successively adding spatial, point-anchor and regional correspondence costs to the connectivity cost, evaluating each fit on the 19-pair benchmark (Methods; Fig. 2a). Using only the Gromov–Wasserstein connectivity cost^25^ gave an AUROC of 0.69, top-1 target-region recovery of 0% and mean centroid displacement of 29 mm. Adding the spatial cost derived from the point-anchor transformation increased AUROC to 0.97 and top-1 recovery to 27%, while reducing displacement to 11 mm. Adding a direct cost for the same point anchors gave an AUROC of 0.93 and top-1 recovery of 26%. Adding the 26 regional correspondence constraints then increased top-1 recovery to 57% and the fraction of mass received by the expected region to 0.54, and reduced displacement to 8.8 mm, although AUROC decreased to 0.90.

**Figure 2.**
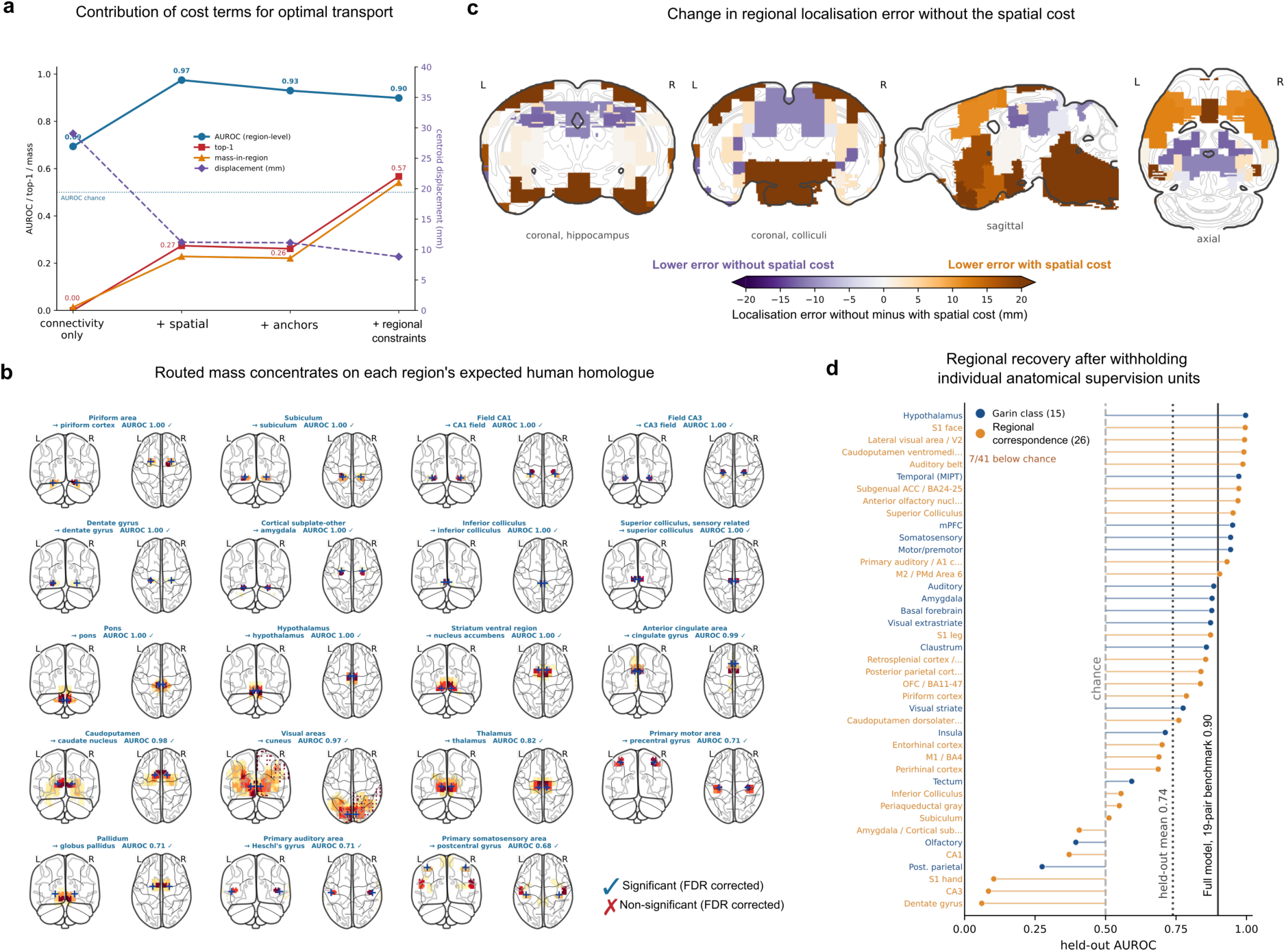
Effects of connectivity, spatial position and anatomical supervision on regional correspondence. **a**, Four cumulative cost configurations evaluated on the 19-pair regional benchmark. Starting with functional and structural connectivity, successive fits added the point-anchor-derived spatial cost, direct point-anchor costs and regional correspondence constraints. The left axis shows area under the receiver operating characteristic curve (AUROC; blue circles), top-1 target-region recovery (red squares) and the fraction of transported mass received by the expected region (orange triangles). The right axis shows mean centroid displacement in millimetres (purple diamonds). **b**, Full-model human correspondence maps for the 19 mouse-region queries, shown as coronal (left) and axial (right) projections. Received weights are summed across source mouse parcels and scaled to each map’s maximum; values below 5% of the maximum are not shown. Blue crosses mark expected human-region reference locations. Labels give the region pair and AUROC; ticks indicate transported-mass enrichment under uniform sampling of human parcels with replacement (false-discovery-rate q < 0.05 across 19 pairs). **c**, Change in localisation error after removing the spatial cost while retaining connectivity, shown on mouse sections. For each benchmark target, all overlapping anatomical constraints were withheld before refitting. Error is the distance between the mass-weighted human prediction centroid and the reference human-region centroid. Colours show error without minus error with the spatial cost: brown indicates lower error with the spatial cost, purple lower error without it. Colours are clipped at ±20 mm (range, −29.9 to +33.7 mm). The three-configuration comparison is shown in Extended Data Fig. 4. **d**, Recovery after omitting each of 41 anatomical supervision units in turn, retaining connectivity and spatial costs. Points show AUROC for the omitted unit (blue, 15 Garin anatomical classes; orange, 26 regional correspondences); horizontal segments connect each point to 0.5. Vertical lines mark chance AUROC (dashed, 0.5), the unweighted mean across 41 omitted units (dotted, 0.74) and the full-model reference from the separate 19-pair benchmark (solid, 0.90).

In the full model, transported mass was enriched in the expected human region for all 19 benchmark pairs under a null based on uniform sampling of human parcels with replacement (FDR q < 0.05; Fig. 2b). Localisation error was lower than expected under the target-centroid rotation null for 16 of 19 pairs (FDR q < 0.05; Extended Data Fig. 4a).

Because some benchmark regions overlapped the anatomical supervision, we refitted OTTER after withholding all point-anchor and regional constraints overlapping each target in turn. Any affected spatial transformation was refitted from the remaining point anchors. With the selected hyperparameters fixed, we compared refits containing both connectivity and spatial costs, or omitting either cost; all retained anatomical supervision elsewhere. Mean localisation errors were 17.9 mm with both costs, 18.2 mm without connectivity and 24.2 mm without the spatial cost. These configurations gave the lowest error in ten, three and six regions, respectively (Fig. 2c; Extended Data Fig. 4b,d). Removing the spatial cost reduced error in the superior colliculus (34.7 to 4.8 mm), thalamus (16.1 to 6.9 mm) and CA1 (23.9 to 11.5 mm). Thus, the lowest mean error was obtained with both costs, although omitting the spatial cost improved localisation in six regions.

In a separate analysis, we assessed recovery after withholding individual anatomical supervision units. We refitted OTTER after omitting each of 41 units in turn: 15 Garin anatomical classes and 26 curated regional correspondences. Each refit retained the connectivity and spatial costs and the remaining anatomical supervision; omitted point anchors were also excluded when refitting the spatial transformation (Methods). Mean AUROC across the 41 omitted units was 0.74, with means of 0.79 for Garin classes and 0.71 for regional correspondences. Seven units had AUROCs below 0.5 (Fig. 2d). The fraction of mouse parcels within each omitted unit whose highest-weight human target lay in the expected region (top-1 recovery) averaged 9.6% across the 41 units. The lowest AUROCs were observed for hippocampal subfields, a somatotopic correspondence in primary somatosensory cortex, and the posterior parietal and olfactory classes. Recovery therefore varied across omitted units under the remaining connectivity, spatial and anatomical guidance, with limited top-1 target-region recovery.

### Cross-species correspondence in functional, microstructural and molecular brain maps

We next assessed whether mouse maps translated through the fitted coupling agreed with human reference maps. These comparisons covered functional networks, functional-connectivity gradients, cortical microstructure and marker-gene expression (Fig. 3).

**Fig. 3.**
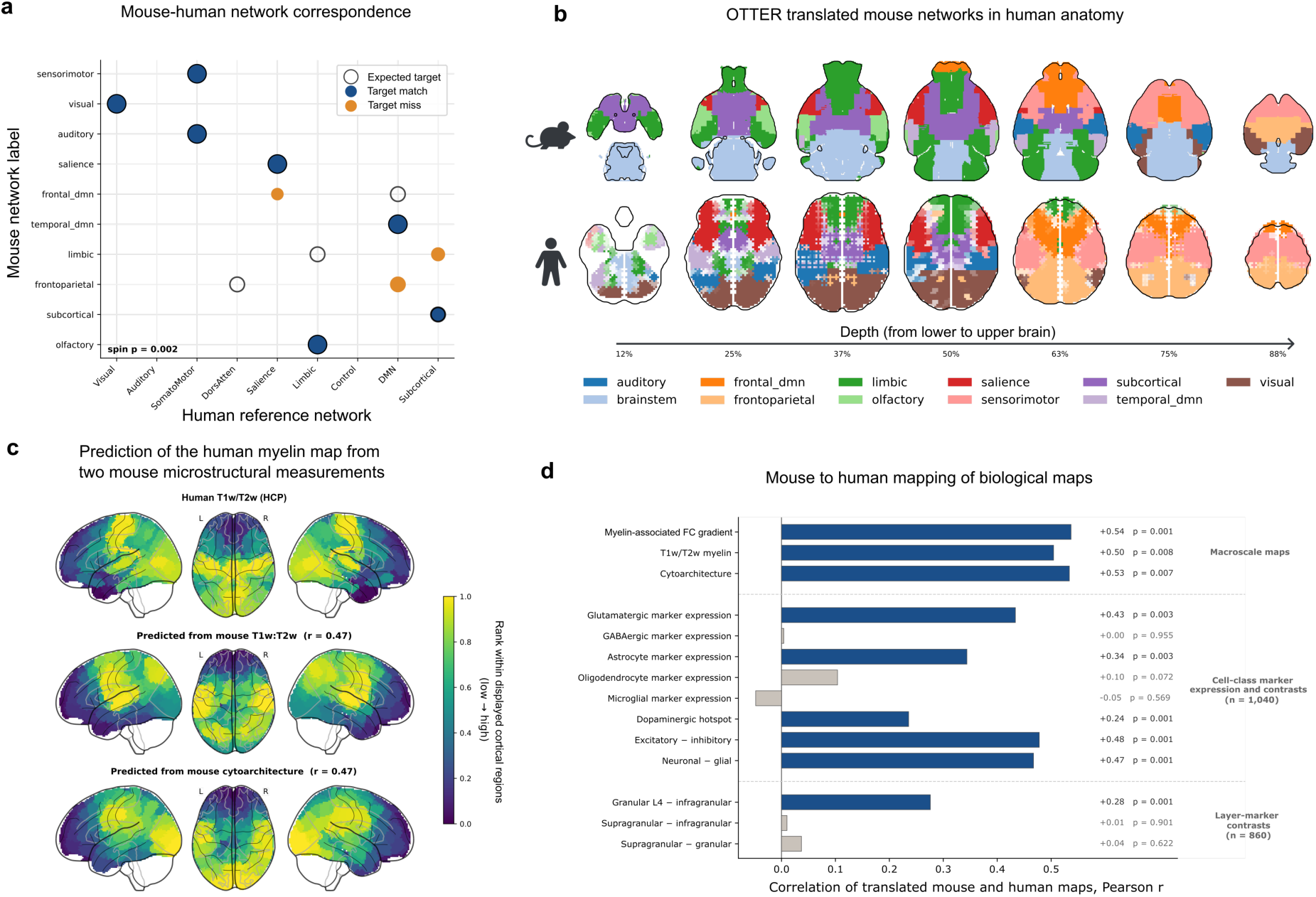
Cross-species correspondence in functional, microstructural and marker-expression maps. **a**, Mouse network labels compared with human reference networks, using cortical labels based on Yeo et al.^31^ (Methods). Mouse labels were assigned to anatomical classes that also guided OTTER’s fit. Open circles indicate expected counterparts; filled circles show the highest-scoring human target for each mouse network (blue, expected match; orange, different target) and circle area scales by total received mass. Seven of ten evaluated networks matched their expected counterpart (one-sided source-label 500 rotation spin test P = 0.002). **b**, Assignments of mouse parcels to eleven networks (top) and translated assignments in human anatomy (bottom). Each human parcel is coloured by the mouse network contributing the greatest total coupling weight. Fainter human colours indicate a smaller difference between the two largest network contributions, relative to total received weight. Axial sections progress from inferior to superior; percentages indicate position within each species’ displayed extent. **c**, Human T1w/T2w myelin proxy (top) and maps translated from mouse T1w/T2w ratios (middle) and ordinal cytoarchitectural types (bottom). Colours show each map’s rank within the 388 represented Schaefer regions. Both translated maps correlated with the human reference at Pearson r = 0.47 across these regions. **d**, Pearson correlations between fourteen translated mouse maps and their human comparison maps: three macroscale maps, eight cell-class marker measures and contrasts, and three layer-marker contrasts. The macroscale comparisons used 2,094, 1,789 and 1,787 human parcels, respectively, in row order. Cell-class comparisons used 1,040 human parcels containing donor tissue; layer contrasts used the 860 cortical parcels within this set. Blue and grey indicate unadjusted two-sided translation-spin P < 0.05 and P ≥ 0.05, respectively, using 1,000 rotations of each mouse input followed by translation through the fixed coupling. Cell-class and layer measures quantify regional marker-gene expression.

We first assessed network-level consistency using eleven curated mouse network labels assigned to the Garin anatomical classes^23^ (note that these anatomical classes also guided OTTER’s optimisation). After translation, the expected human network received the most coupling mass for seven of the ten mouse networks including, for example, the sensorimotor, visual and subcortical networks (Fig. 3a). This count was greater than expected under the source-label spin null (P = 0.002; 500 rotations). The translated network territories are shown in Fig. 3b. Separately, the highest-weight human targets of mouse parcels were less dispersed around their network’s target centroid than the mean under coupling-row permutations in nine of the eleven networks (Methods).

We also derived seven mouse network maps by applying independent component analysis (ICA) to the functional-connectivity matrix used to fit OTTER. Six components had dominant anatomical network labels with specified human counterparts and were evaluated (Methods). After translation, the sensorimotor, salience and limbic components had their highest mean values in the expected human networks, whereas frontoparietal, frontal default-mode and temporal default-mode components did not.

We next assessed translation of functional-connectivity gradients, which describe continuous variation in connectivity patterns^27^. We used a diffusion-map embedding to derive the first four functional-connectivity gradients separately in mouse and human and selected the component with the largest absolute Spearman correlation with that species’ T1w/T2w ratio map. This myelin-associated gradient was the second component in both species (Spearman |ρ| = 0.56 in mouse and 0.59 in human). After translation, the mouse gradient correlated with the selected human gradient at |r| = 0.54. The third mouse and human components also showed cross-species correspondence (|r| = 0.65). After refitting OTTER without the connectivity cost, the corresponding correlations were |r| = 0.47 and 0.63, respectively (Extended Data Fig. 6).

We then assessed cross-species correspondence in cortical microstructure using two mouse maps from Fulcher et al.^28^: T1w/T2w ratios and ordinal cytoarchitectural types (an ordered classification of cortical regions based on how distinctly their cell layers are organised). Both maps were translated through the fitted coupling and compared with the human T1w/T2w myelin proxy^29^. The translated T1w/T2w map correlated with the human map at |r| = 0.50 across 1,789 human parcels, and the translated cytoarchitectural-type map at |r| = 0.53 across 1,787 human parcels (unadjusted two-sided translation-spin P = 0.008 and 0.007, respectively; Fig. 3d and Extended Data Fig. 7). At the regional level, both correlations were r = 0.47 across 388 represented Schaefer atlas regions (Fig. 3c).

We next tested correspondence between translated mouse marker-expression maps and their human counterparts, using eight cell-class measures and three layer-marker contrasts (Fig. 3d). Human comparisons were restricted to parcels containing Allen Human Brain Atlas donor tissue: 1,040 parcels for the cell-class measures and 860 cortical parcels for the layer contrasts. Six of the eleven measures showed significant positive correspondence: five cell-class measures and the granular layer 4–infragranular contrast (r = 0.24–0.48; unadjusted translation-spin P = 0.001–0.003). GABAergic, oligodendrocyte and microglial marker expression and the supragranular–infragranular and supragranular–granular contrasts did not show significant correspondence (|r| ≤ 0.11; unadjusted P = 0.072–0.955). Using a separate human cortical mask, the four cell-class measures examined had weaker correlations across 856 parcels containing donor tissue (r = 0.12–0.25; Methods). Together, the map comparisons identified cross-species correspondence in selected functional, microstructural and marker-expression maps.

### Regional correspondence and phenotype translation compared with TransBrain

To compare regional correspondence, we evaluated OTTER and TransBrain^16^ on 24 mouse-region queries over a common set of 127 human target regions, using reference correspondences supplied with the TransBrain package. OTTER’s coupling weights were summed over parcels within each source mouse region and each human target region in the Brainnetome-based configuration (Methods). Mean AUROC across queries was 0.83 for OTTER and 0.84 for TransBrain (paired Wilcoxon signed-rank test, P = 0.36; Fig. 4a). OTTER was not fitted to TransBrain outputs or transcriptomic embeddings, however, some queried mouse regions overlapped OTTER’s regional correspondence constraints.

**Fig. 4.**
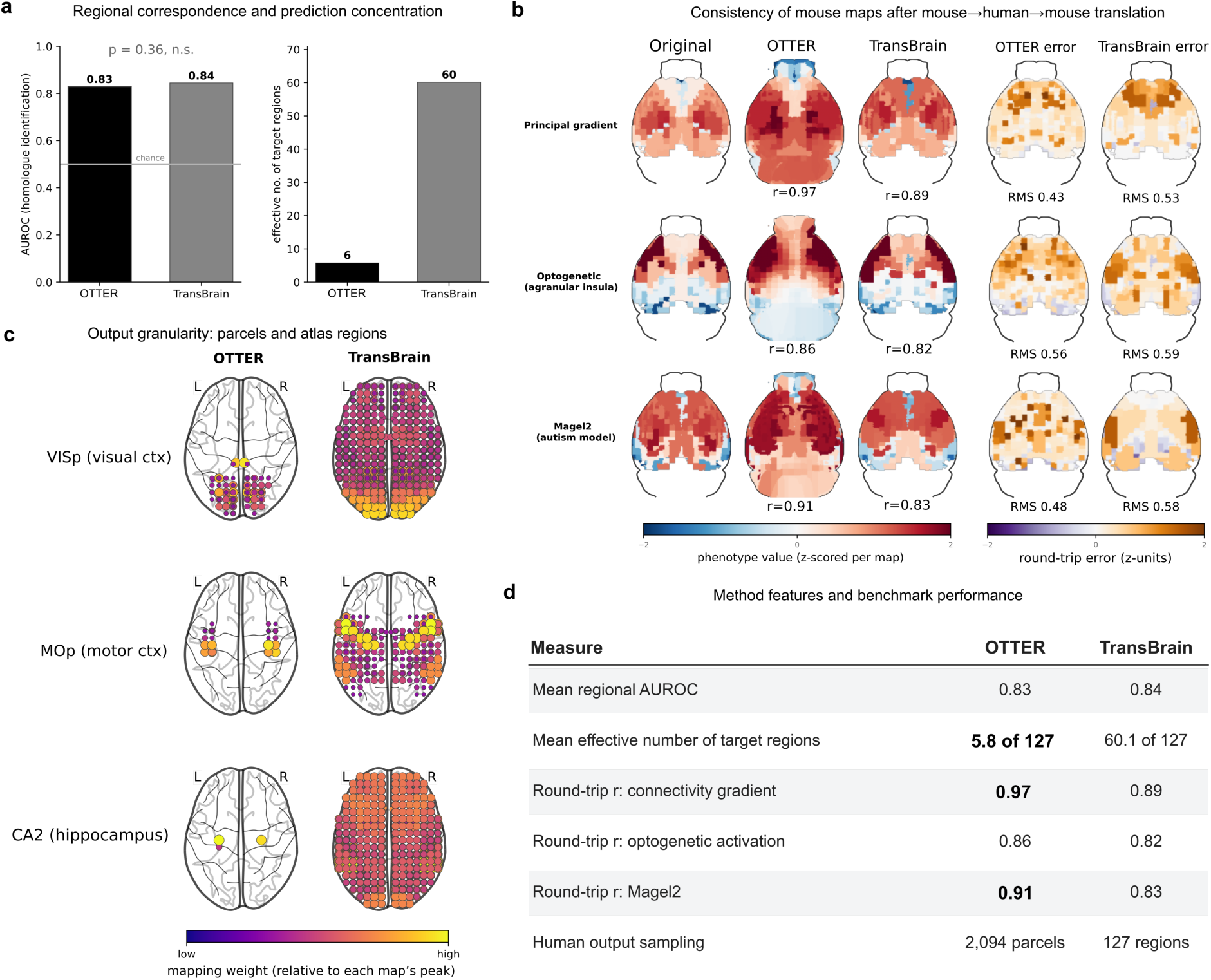
Comparison of OTTER and TransBrain for regional correspondence and map translation. **a**, Mean area under the receiver operating characteristic curve (AUROC; left) and effective target count (right) across 24 mouse-region queries over the same 127 human target regions, using reference correspondences from the TransBrain repository. Mean AUROCs were 0.83 and 0.84 for OTTER and TransBrain, respectively (two-sided paired Wilcoxon signed-rank test, P = 0.36; n = 24 queries). Effective counts were calculated as the inverse sum of squared normalised regional weights and averaged across queries (approximately 6 and 60, respectively). Some queried mouse regions overlapped OTTER’s regional correspondence constraints on the mouse side, so this evaluation was not fully held out. **b**, Original mouse maps, maps returned after mouse→human→mouse translation using OTTER and TransBrain, and their respective errors (left to right). Rows show a functional-connectivity gradient, fMRI activation following optogenetic stimulation of the insula and standardised regional volume differences between Magel2-mutant and control mice. Pearson r values compare original and returned regional values over the same 52 mouse regions with measured inputs. Within each row, maps were separately z-scored on their common finite parcel mask. Errors are returned minus original values; RMS denotes root-mean-square error across these parcels. Original and TransBrain maps are constant within regions; OTTER maps retain within-region variation. **c**, Human outputs for mouse primary visual cortex (VISp), primary motor cortex (MOp) and hippocampal area CA2. OTTER (left; 2,094 parcels) and TransBrain (right; 127 atlas regions) are displayed at the same human parcel positions, repeating each TransBrain regional value at its constituent parcels. Colour and marker size indicate weight relative to each map’s peak; only weights above 5% of the peak are shown. **d**, Descriptive summary of numerical comparisons and output properties for the two methods.

On the same 127 human target regions, both methods ranked a reference human region first in 25% of the 24 mouse queries. Top-3 recovery was 46% for OTTER and 50% for TransBrain, and top-5 recovery was 67% and 58%, respectively. The mean fraction of normalised weight assigned to reference human regions was 0.21 for OTTER and 0.07 for TransBrain. The mean effective number of target regions for these normalised weights was approximately 6 for OTTER and 60 for TransBrain, indicating more concentrated OTTER predictions (Fig. 4a).

To assess round-trip consistency, we translated three regional mouse maps to human space and back: a functional-connectivity gradient, an insula optogenetic activation map and the Magel2 map (a map of standardised regional volume differences between Magel2-mutant and control mice). Both methods were scored against the original regional values on the same 52 mouse regions with measured inputs. Pearson correlations for OTTER and TransBrain, respectively, were 0.97 and 0.89 for the gradient, 0.86 and 0.82 for the insula map, and 0.91 and 0.83 for the Magel2 map (Fig. 4b). In a separate forward-gradient comparison across 101 human cortical regions, absolute Pearson correlations with the observed human gradient were 0.56 for OTTER and 0.52 for TransBrain. The absolute Pearson correlation between the two predictions was 0.83.

OTTER provided correspondence weights for 2,094 human parcels, whereas the TransBrain configuration evaluated here provided values for 127 atlas regions. To illustrate this difference, we displayed both outputs on the same human parcel grid, assigning the same TransBrain regional value to every parcel within that region. The examples for mouse primary visual cortex, primary motor cortex and hippocampal area CA2 show variation in OTTER’s predicted correspondence within the larger atlas regions (Fig. 4c). Figure 4d summarises the numerical comparisons and output properties of the two configurations.

### Mouse-based reconstruction of human functional connectivity is less accurate in association cortex

Anatomically corresponding regions can differ in their connectivity to the rest of the brain. To characterise these differences, we reconstructed group-mean human functional connectivity from the group-mean mouse connectome using the fitted OTTER coupling. For each human parcel receiving coupling weight, we normalised its mouse correspondence weights to sum to one. We then estimated connectivity between each pair of human parcels as the weighted average of mouse connectivity between their corresponding mouse parcels *F^_H_ = AF_M_A^T^* where *F*_M_ is the measured mouse functional-connectivity matrix and *A* contains the normalised weights, with human parcels in rows and mouse parcels in columns. For each human parcel, we calculated the Pearson correlation between its reconstructed and measured connectivity profiles across all other human parcels (Methods). Higher scores indicate closer agreement in which human targets are relatively strongly or weakly connected to that parcel.

The highest reconstruction score for an individual parcel was found in the Visual network (r = 0.77; Fig 5a top), whereas the lowest-scoring parcel was found in the Control B network in the left dorsal frontal cortex (r = 0.07; Fig. 5a bottom). Across 1,824 scored cortical parcels, reconstruction scores averaged 0.45 and ranged from −0.13 to 0.77. Scores correlated strongly between matched left and right parcels (Spearman ρ = 0.86) and between regional mean scores in matched Schaefer atlas regions (ρ = 0.61). The cortical map showed higher scores in sensorimotor, auditory and visual territories and lower scores in prefrontal and lateral temporal cortex (Fig. 5b).

**Fig. 5.**
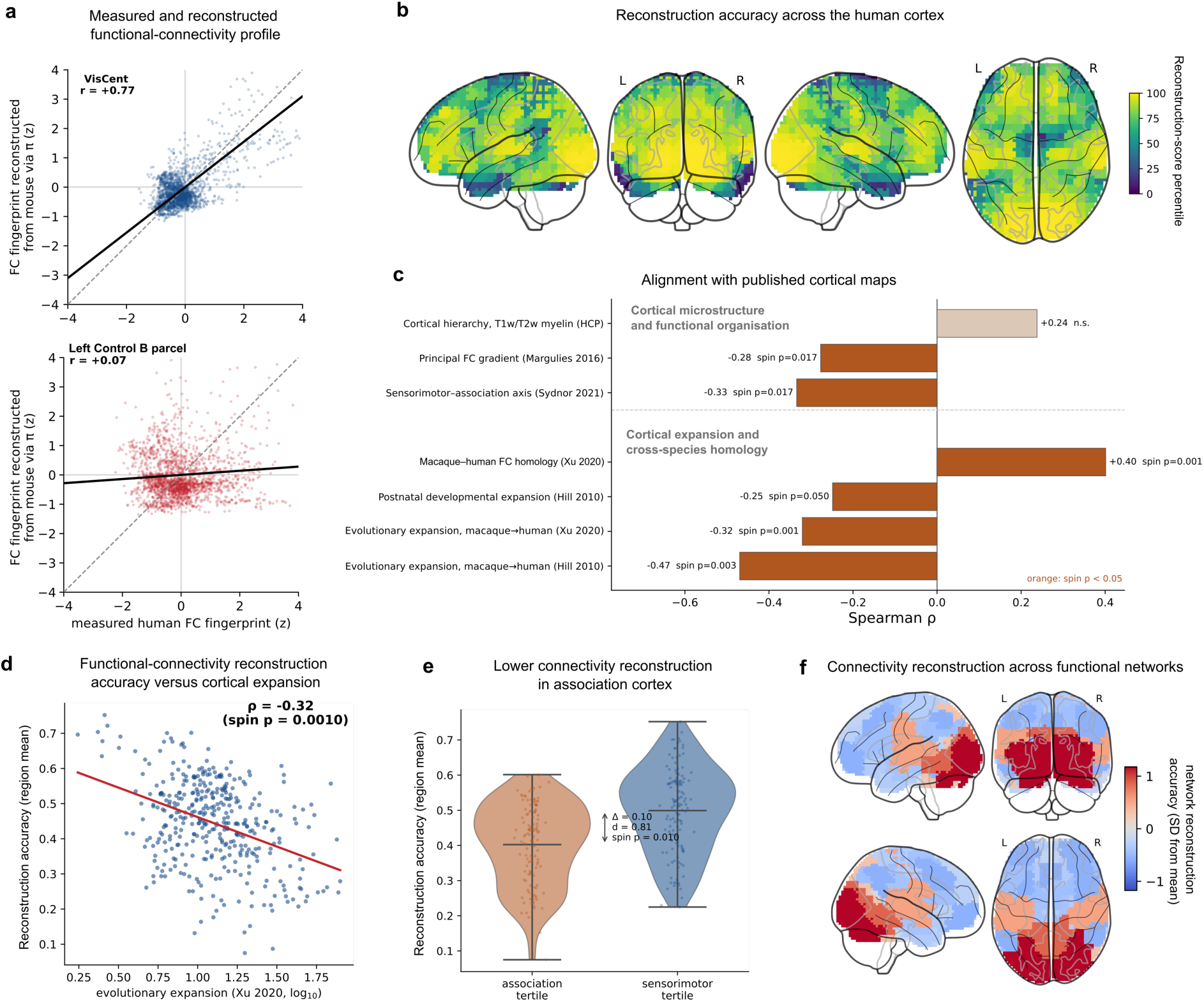
Mouse-based reconstruction of human functional connectivity is less accurate in expanded association cortex. Reconstruction scores are Pearson correlations between reconstructed and measured group-mean connectivity profiles for each human parcel. The connectomes also contributed to fitting OTTER, so this is an internal assessment. **a**, Measured versus reconstructed connectivity for a visual-network parcel (top, r = 0.77) and a left Control B network parcel (bottom, r = 0.07). Each point represents connectivity to another human parcel; profiles are separately z-scored. Solid black lines show least-squares fits; dashed grey lines indicate identity. **b**, Reconstruction scores expressed as percentile ranks across the 1,824 scored cortical parcels. **c**, Spearman correlations between mean reconstruction scores within Schaefer regions and seven published cortical maps (197 right-hemisphere regions for each of the two Hill expansion maps; 391 regions for each other comparison). The MRI measure was the T1w/T2w myelin proxy. Orange and tan indicate unadjusted two-sided cortical-map spin P < 0.05 and P ≥ 0.05, respectively (1,000 rotations). The Xu expansion and functional-homology maps^14^ compare macaques with humans. The reference maps describe overlapping aspects of cortical organisation. **d**, Mean regional reconstruction scores versus macaque-to-human cortical expansion^14^, with expansion displayed on a log₁₀ scale (391 Schaefer regions; Spearman ρ = −0.32, cortical-map spin P = 0.001). The red line is a linear fit to the displayed coordinates. **e**, Regional reconstruction scores in the highest (association) and lowest (sensorimotor) thirds of the sensorimotor–association axis (130 regions per group). Points show regional means; violin widths show density, central horizontal lines show group means, and capped vertical lines span minima to maxima. Group means were 0.40 and 0.50, respectively (sensorimotor minus association, Δ = 0.10; Cohen’s d = 0.81; two-sided cortical-map spin P = 0.010, 1,000 rotations). **f**, Mean reconstruction scores across 16 represented network groups based on the Yeo 17-network parcellation³¹. Colours indicate each network’s mean relative to the mean and standard deviation of all scored cortical parcels.

To relate this spatial pattern to cortical organisation, we averaged reconstruction scores within Schaefer regions and compared them with seven published reference maps (Fig. 5c). Scores were lower in regions with greater macaque-to-human cortical expansion (Hill et al. ^32^, Spearman ρ = −0.47, P = 0.003; Xu et al. ^14^, ρ = −0.32, P = 0.001; Fig. 5d). Scores were also lower towards the association end of the sensorimotor–association axis^30^ (ρ = −0.33, P = 0.017) and the published principal functional-connectivity gradient^27^ (ρ = −0.28, P = 0.017). Conversely, scores were higher where macaque–human functional-connectivity homology^14^ was greater (ρ = 0.40, P = 0.001). Associations were weaker with human postnatal cortical expansion^32^ (ρ = −0.25, P = 0.050) and the MRI-derived T1w/T2w myelin proxy^29^ (ρ = 0.24, P = 0.106). These associations link lower reconstruction scores to expanded association territories.

To quantify differences across the sensorimotor–association axis, we ranked Schaefer regions by their axis values and compared the highest and lowest thirds. Mean regional reconstruction scores were 0.40 in the association tertile and 0.50 in the sensorimotor tertile (sensorimotor minus association, Δ = 0.10; Cohen’s d = 0.81; cortical-map spin P = 0.010; Fig. 5e). The distributions overlapped: 21% of association regions scored above the sensorimotor median. The lowest regional means were 0.07 and 0.22 in the association and sensorimotor tertiles, respectively.

We also compared mean reconstruction scores across the 16 represented network groups based on the Yeo 17-network parcellation^31^. Control B, a subdivision of the frontoparietal control network, had the lowest mean score, followed by Control A and Salience/Ventral Attention A (Fig. 5f). Across the 1,824 scored cortical parcels, the Control B-versus-rest contrast was −0.61 SD, using the standard deviation of reconstruction scores across this parcel set (unadjusted two-sided cortical-map spin P = 0.0105; 2,000 rotations). Note that this contrast did not survive Bonferroni correction across the 16 network tests (adjusted P = 0.168).

To assess whether the regional pattern also occurred in structural connectivity, we reconstructed human structural connectivity from the mouse structural connectome through the unchanged OTTER coupling. Structural reconstruction scores showed little association with macaque-to-human cortical expansion (Spearman ρ = −0.05), and the Control B-versus-rest contrast was +0.19 SD (Extended Data Fig. 5a).

We also assessed cross-species expression similarity across 51 one-to-one orthologous genes in 1,040 human parcels with complete expression profiles and finite reconstruction scores. For each human parcel, we retained the highest cosine similarity between its centred expression profile and that of any eligible mouse parcel (Methods). This expression-based match was selected without the OTTER coupling, and these expression data were not used to fit OTTER. In Control B, best-match expression similarity was 0.18 SD lower than in the other parcels in this set, using the standard deviation of similarity scores across all 1,040 parcels (two-sided spin P = 0.39).

### Bidirectional translation of activation and molecular-imaging maps

To assess whether translated maps localised to expected brain regions, we used the same fitted OTTER coupling for translation in both directions. We translated a mouse experimental activation map into a predicted human map, and human meta-analytic activation and molecular-imaging maps into predicted mouse maps (Fig. 6; Methods).

**Fig. 6.**
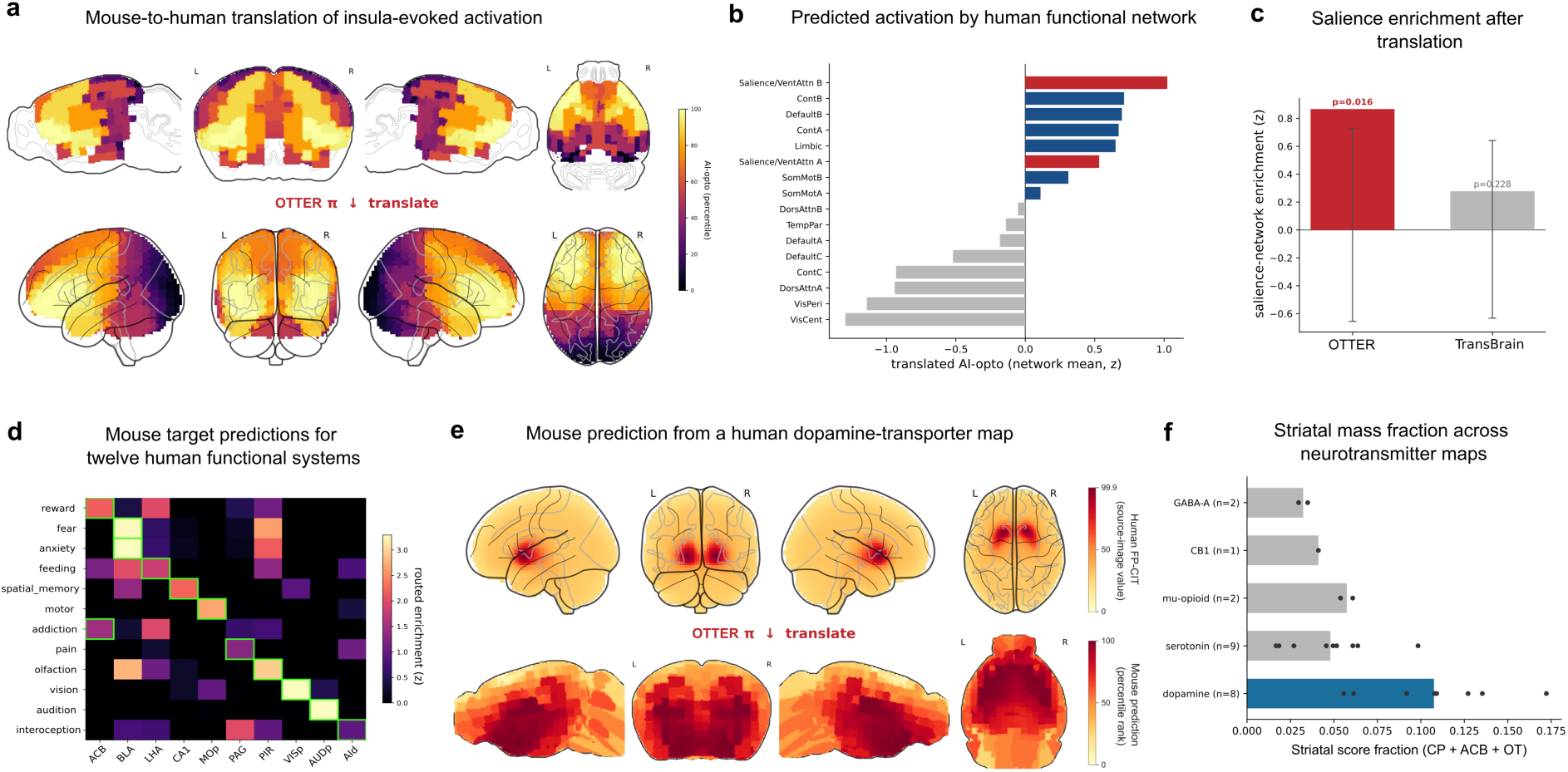
Bidirectional translation of activation and molecular-imaging maps. All OTTER translations used the same fitted coupling. a, Mouse activation following optogenetic stimulation of agranular insula^33^ (top) and its human prediction (bottom), displayed as percentile ranks within each map. b, Mean predicted human activation across 16 represented Yeo-17 network groups, after z-standardisation over all 2,059 human parcels with finite predictions. Red highlights salience/ventral-attention groups; blue and grey show other positive and negative means, respectively. Salience/Ventral Attention B had the highest mean (z = 1.02), and Visual Central the lowest (z = −1.30). c, Mean predicted activation in Salience/Ventral Attention A and B parcels minus that in the remaining parcels, for OTTER (red, 0.87 z units) and TransBrain (grey, 0.28 z units). Each predicted map was z-standardised separately across the same 1,635 eligible human parcels (Methods). Thin capped lines show each shuffled-input null’s mean ± 1.96 standard deviations. Activation values were permuted among mouse input regions before retranslation (1,000 permutations per method; one-sided P = 0.016 and 0.228, respectively). Each P value compares a method with its own null. d, Mouse predictions from Neurosynth meta-analytic maps^34^ associated with 12 functions and behaviours. Colours show mean predicted values standardised across 83 eligible mouse structures within each row; selected expected-target structures are displayed. Green outlines mark one eligible expected target per map (Supplementary Table 2). Nine maps placed at least one expected target among the top three of all 83 structures, each represented by at least five parcels. e, Human FP-CIT dopamine-transporter SPECT input map (neuromaps annotation dukart2018/fpcit; top) and predicted mouse map (bottom). Human colours show rescaled source-image values. Mouse colours show percentile ranks of translated values. f, Sum of positive translated structure means in the caudoputamen, nucleus accumbens and olfactory tubercle divided by the corresponding sum across all 83 eligible mouse structures. Points represent individual molecular-imaging maps; bars show group means. Fractions were higher for eight dopamine maps than for 14 pooled reference maps: serotonin (n = 9), μ-opioid (n = 2), CB1 (n = 1) and GABA-A (n = 2) (exact one-sided Mann–Whitney U = 105, P = 1.4 × 10⁻⁴).

For mouse-to-human translation, we used a whole-brain activation map obtained after optogenetic stimulation of the mouse agranular insula^33^. The predicted human map exhibited high values in the insular and ventral-attention cortex (Fig. 6a). After z-standardisation across the 2,059 human parcels with finite predictions, Salience/Ventral Attention B had the highest mean value among the 16 represented Yeo-17 network labels (mean z = 1.02), whereas Visual Central had the lowest (mean z = −1.30; Fig. 6b). Mean predicted activation in Salience/Ventral Attention A and B parcels exceeded that in the remaining 1,798 human parcels by 0.86 z units (one-sided coupling-row permutation P = 0.001).

To compare this network enrichment with TransBrain, we z-standardised each predicted map across the same 1,635 eligible human parcels and subtracted the mean for the remaining parcels from the mean for Salience/Ventral Attention A and B parcels. Enrichment was 0.87 z units for OTTER and 0.28 for TransBrain (Fig. 6c). We assessed each result by shuffling activation values among the mouse input regions and repeating the translation (1,000 permutations per method; Methods). The corresponding one-sided permutation P-values were 0.016 for OTTER and 0.228 for TransBrain, each relative to its own shuffled-input distribution.

We next assessed human-to-mouse translation using Neurosynth meta-analytic activation maps associated with 12 functions and behaviours, including vision, movement, fear and reward^34^. For each translated map, we ranked 83 mouse structures, each represented by at least five parcels, by their mean predicted value. Expected mouse targets were hypothesised before scoring (Supplementary Table 2; Methods). Nine of the 12 maps placed at least one expected target among the top three structures, and three placed an expected target first (Fig. 6d). The feeding, addiction and pain maps did not place an expected target among the top three. The visual and motor maps gave high predicted values in primary visual and motor cortex, respectively. Separately, we tested the highest score among each map’s eligible expected targets against the same statistic after 1,000 rotations of the human source map and retranslation through the fixed coupling. All 12 maps showed significant expected-target scores in this test (unadjusted one-sided P = 0.001–0.033).

Finally, we assessed reverse translation of molecular maps using the dopaminergic system, which is disrupted in Parkinson disease^35^. We translated eight human maps covering D1 and D2/3 receptor binding and presynaptic dopamine transporters^35–37^. For all eight maps, the maximum of the mean predicted values in caudoputamen, nucleus accumbens and substantia nigra pars reticulata was significant relative to the reverse-translation spin null (unadjusted one-sided P ≤ 0.005; Fig. 6e). We summarised each prediction by dividing the sum of positive translated structure means in caudoputamen, nucleus accumbens and olfactory tubercle by the corresponding sum across all 83 eligible mouse structures (Methods). These fractions were higher for the eight dopamine maps than for the 14 pooled serotonin, μ-opioid, cannabinoid CB1 and GABA-A reference maps (exact one-sided Mann–Whitney U = 105, P = 1.4 × 10⁻⁴; Fig. 6f) ^37–39^.

For the human FP-CIT dopamine-transporter map, mean input values correlated with mean values in the translated mouse prediction across the 21 paired anatomical classes defined by each parcel’s nearest same-hemisphere point anchor (Spearman ρ = 0.84, two-sided asymptotic P = 1.9 × 10⁻⁶; Extended Data Fig. 8a). This comparison assessed preservation of the class-level ranking in the derived mouse prediction. Together, the activation and molecular-map analyses assessed translation against expected regional patterns in both directions. We next examined maps associated with human disease phenotypes, symptoms and stages.

### Mouse regional predictions from human phenotype and symptom maps

Clinical presentations of the same disease can involve different brain regions. We therefore examined which mouse brain regions were prioritised when OTTER translated human maps associated with disease phenotype, symptom or stage. We began with Alzheimer disease (AD), using published tau positron emission tomography (tau-PET) and voxel-based morphometry contrast maps derived from 119 amyloid-positive participants^40^. The groups comprised posterior cortical atrophy (PCA), logopenic variant primary progressive aphasia (lvPPA) and AD without either syndrome. PCA and lvPPA primarily affect visuospatial and language function, respectively. Each group was contrasted with the other two; positive T values indicated greater tau binding or lower grey-matter volume in the named phenotype (Methods).

Before scoring, we assigned each phenotype to a candidate mouse target set: the entorhinal cortex and hippocampus for AD without PCA or lvPPA; the visual cortex for PCA; and the auditory and temporal association cortices for lvPPA^41–44^. For each phenotype map, we defined selectivity as the correspondence-weighted score for the assigned target set minus the mean score for the other two sets (Methods) and joint selectivity (C) was the sum of these three differences. All three phenotype selectivities were positive for both modalities, and joint selectivity was greater than expected under 10,000 spatial rotations (tau-PET: C = 2.565, P = 0.0046; voxel-based morphometry: C = 2.354, P = 0.0046; one-sided tests; Fig. 7a,b).

**Fig. 7.**
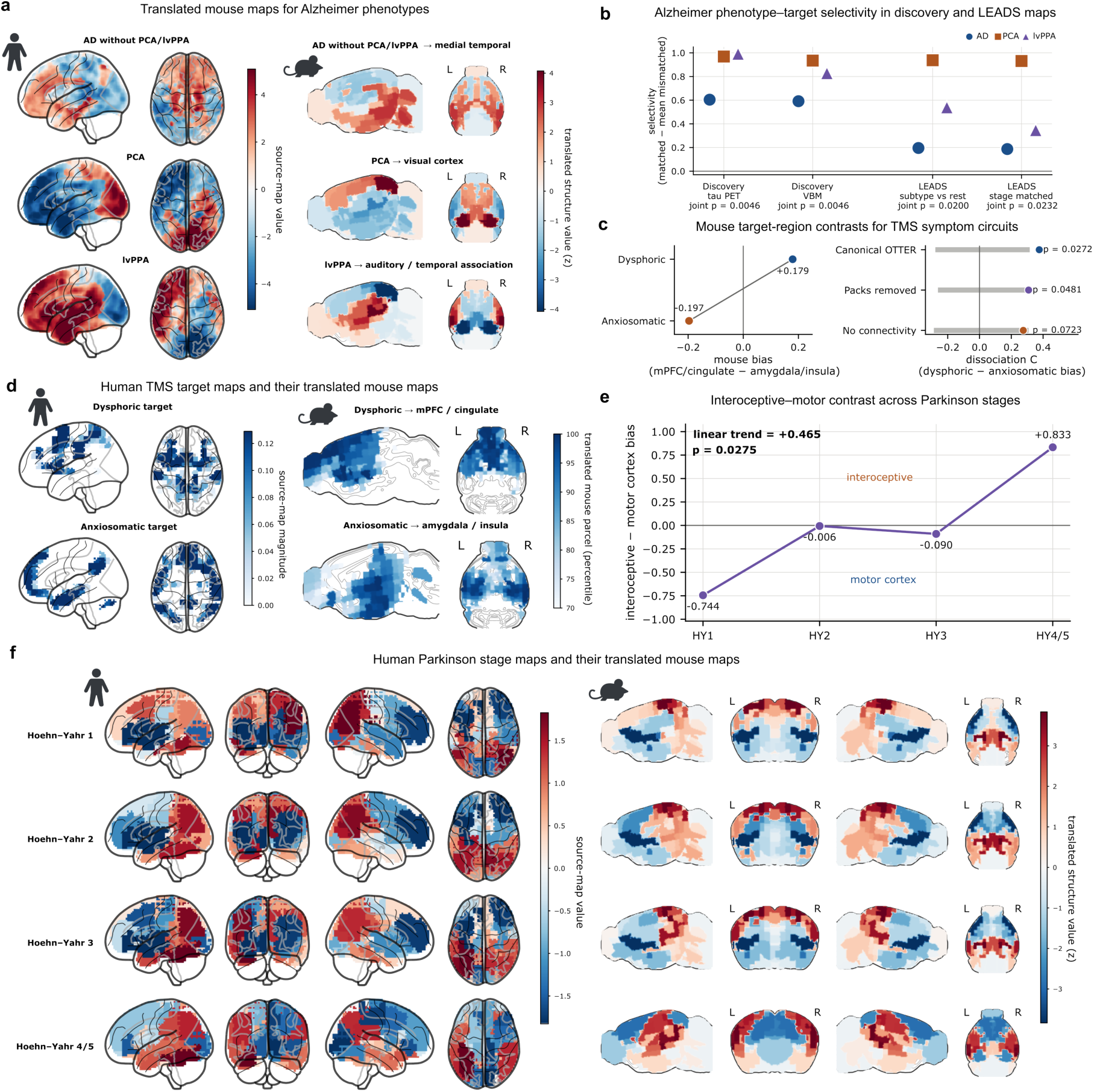
Mouse regional predictions from human disease phenotype, symptom and stage maps. Translations used full OTTER unless otherwise indicated. Mouse maps in a and f show translated structure means z-standardised across structures within each map. **a**, Human tau-PET contrasts (left), each clinical group versus the other two^40^, and corresponding mouse predictions (right). Groups were Alzheimer disease (AD) without posterior cortical atrophy (PCA) or logopenic variant primary progressive aphasia (lvPPA), PCA and lvPPA. **b**, Target selectivity for discovery tau-PET and voxel-based morphometry, Longitudinal Early-Onset Alzheimer’s Disease Study (LEADS) subtype-versus-rest maps and unthresholded contrasts averaged across matched tau-stage ranges^45^. Circles, squares and triangles denote entorhinal/hippocampal, visual and auditory/temporal-association target sets, respectively. These were assigned to AD, PCA and lvPPA in discovery, and to typical, posterior and left-temporal tau subtypes in LEADS. Selectivity is the assigned-set score minus the mean score for the other two sets. Displayed P values test the sum of three selectivities (one-sided tests; 10,000 spatial rotations). Each point represents an assignment. **c**, Mouse regional bias for dysphoric and anxiosomatic transcranial magnetic stimulation (TMS) inputs (left), defined as medial prefrontal/cingulate minus amygdala/insula scores. The between-input contrast C is dysphoric minus anxiosomatic bias. Right, observed C values and 95% spatial-null ranges for full OTTER, the refit without the amygdala, cingulate and lateral-prefrontal regional constraint sets, and the matched model without the connectivity cost. At the primary 10% threshold, values were C = 0.376, 0.309 and 0.275 (corrected P = 0.0272, 0.0481 and 0.0723), respectively. The paired full-minus-no-connectivity contrast was ΔC = 0.100 (corrected P = 0.0311). One-sided tests used 10,000 Moran surrogates and maximum-statistic family-wise-error correction across four input-tail thresholds (5%, 10%, 20% and 30%). **d**, Human TMS targeting-atlas inputs^46^ (left), using the primary 10% negative (dysphoric) and positive (anxiosomatic) tails, and mouse predictions (right). Human colours show input magnitudes; mouse colours show parcel percentile ranks, displaying values at or above the 70th percentile. **e**, Interoceptive-minus-motor contrast for each cross-sectional Parkinson stage-group map. The interoceptive set comprised visceral, gustatory and agranular-insular regions; the motor set comprised primary and secondary motor cortex. The contrast was −0.744 at stage 1 and +0.833 at stage 4/5 (ordered-group linear trend = 0.465; two-sided spatial-null P = 0.0275, 10,000 rotations). **f**, Human ENIGMA cortical-thickness effects for Hoehn–Yahr stages 1, 2, 3 and 4/5 versus matched controls^48^ (left), and mouse predictions (right). Human effects were sign-reversed and z-standardised within each stage map; higher values indicate relatively greater thinning within that map.

To assess dependence on the corresponding anatomical constraints, we refitted OTTER without the eight directly relevant sets of regional correspondence constraints, retaining the remaining anatomical supervision and spatial transformation. Joint selectivity remained significant for both tau-PET (C = 2.830, P = 0.0047) and voxel-based morphometry (C = 2.674, P = 0.0048; Extended Data Fig. 8b).

To evaluate the same mouse target assignments in an external dataset, we analysed published baseline tau-PET maps from the multisite Longitudinal Early-Onset Alzheimer’s Disease Study (LEADS; 359 participants)^45^. These maps compared each of three tau-defined subtypes (typical, posterior and left-temporal) with the other two. We assigned the typical subtype to entorhinal cortex and hippocampus, the posterior subtype to visual cortex, and the left-temporal subtype to auditory and temporal association cortex (Methods). All three selectivities were positive under full OTTER, and joint selectivity was significant for both the full model (C = 1.673, P = 0.020) and the refit without the eight regional constraint sets (C = 1.928, P = 0.015; one-sided spatial-null tests; Fig. 7b and Extended Data Fig. 8b).

We also constructed contrasts between unthresholded subtype mean maps within ranges of model-estimated tau stage, then averaged these contrasts across ranges (Methods). All three selectivities were again positive under full OTTER, and joint selectivity remained significant for both the full model (C = 1.465, P = 0.023) and the refit (C = 1.690, P = 0.022; one-sided spatial-null tests; Fig. 7b and Extended Data Fig. 8b).

To examine symptom-related differences within depression, we used a published transcranial magnetic stimulation (TMS) targeting atlas derived from symptom responses and functional connectivity^46^. The atlas distinguishes potential targets for dysphoric symptoms (low mood and loss of interest) from those for anxiosomatic symptoms (anxiety and somatic complaints). A subsequent randomised trial found greater anxiety improvement with stimulation at the anxiosomatic target than at the dysphoric target^47^.

We translated the atlas’s negative and positive components as dysphoric and anxiosomatic inputs, respectively, using the primary 10% tail threshold (Methods). For each translated map, we calculated regional bias as the score (sum of translated values across its mouse parcels divided by the sum across all named mouse parcels) in mouse medial prefrontal and cingulate regions minus that in amygdala and insula. C was the dysphoric bias minus the anxiosomatic bias. The dysphoric map favoured medial prefrontal and cingulate regions, whereas the anxiosomatic map favoured amygdala and insula (C = 0.376, P = 0.0272; one-sided spatial-null test, family-wise-error corrected across four tail thresholds; Fig. 7c,d). This separation persisted after refitting OTTER without the amygdala, cingulate and lateral-prefrontal regional correspondence constraint sets, retaining the remaining anatomical guidance (C = 0.309, corrected P = 0.0481; Extended Data Fig. 8c,d).

### Human Parkinson disease stage maps and longitudinal mouse MRI

To examine stage-related differences in Parkinson disease, we translated cross-sectional ENIGMA maps of cortical-thickness differences between people at Hoehn–Yahr stages 1, 2, 3 and 4/5 and age- and sex-matched controls^48^. We standardised each map across regions so that the translated scores represented relative spatial patterns of thinning (Methods). We compared scores in mouse visceral, gustatory and agranular-insular regions (the interoceptive set) with those in primary and secondary motor cortex, defining bias as interoceptive minus motor. This bias was negative at stage 1 (−0.744), close to zero at stages 2 (−0.006) and 3 (−0.090), and positive at stage 4/5 (+0.833), with a positive overall linear trend across the four ordered stage groups (linear trend = 0.465, two-sided spatial-null P = 0.0275; Fig. 7e,f). The stage trend had the same direction in both hemisphere-specific analyses.

To assess this stage-related pattern across individuals in an external cohort, we analysed 141 participants with Parkinson disease and 69 controls from the Quebec Parkinson Network Neuroimaging Cohort (QPN-NC). We used the controls to estimate each patient’s thinning map and scored its spatial pattern using the same OTTER interoceptive-minus-motor contrast (Methods). Across the four ordered Hoehn–Yahr stage groups, higher stage was associated with higher scores after adjustment for MRI age, sex and the interval between clinical assessment and MRI (partial Spearman ρ = 0.164, one-sided spatial-null P = 0.0222; Extended Data Fig. 10a).

An omnibus spatial test across this participant-level association and four models of regional stage effects was significant (maximum-statistic P = 0.0493; Extended Data Fig. 10b). For all four models, the translated regional associations between cortical thinning and stage gave a positive interoceptive-minus-motor contrast (Extended Data Fig. 10b). For the primary regional model, the contrast also remained positive after excluding each participant with Parkinson disease or each control in turn (Extended Data Fig. 10c). Positive regional contrasts were also retained in both hemispheres using the alternative 148-region Destrieux parcellation (Extended Data Fig. 10d).

To assess whether the anatomical contrast identified across human Parkinson disease stages also changed over time in mice, we reanalysed longitudinal structural MRI from wild-type mice and M83 mice expressing human A53T α-synuclein. These mice received right-striatal injections of human or mouse α-synuclein preformed fibrils (PFFs), or phosphate-buffered saline (PBS) ^49^. At each follow-up, we estimated regional PFF-associated atrophy relative to genotype-matched PBS controls, adjusting for baseline MRI measurements and sex. We then spatially standardised each effect map and calculated the mean atrophy effect across visceral, gustatory and agranular-insular regions minus that across primary and secondary motor cortex. Averaged equally across the four genotype-by-PFF conditions, this interoceptive-minus-motor score increased from 30 to 120 days after injection (slope = 0.0192 score units per day, one-sided animal-label permutation P = 0.0065; Extended Data Fig. 11a,b). All four conditions had positive overall slopes, and the mean slope remained positive after adjustment for cohort and acquisition batch, restriction to animals with all three follow-ups, and changes to spatial normalisation (Extended Data Fig. 11c). This agreement in direction with the human stage comparisons motivates experiments examining these interoceptive regions alongside motor cortex in Parkinson disease models^50^.

Together, these applications show how OTTER uses human imaging to identify mouse brain regions for experiments tailored to disease phenotype, symptom and stage.

## Discussion

Here, we have introduced OTTER, a probabilistic framework for translating experimental and clinical brain maps between mouse and human. By combining functional and structural connectivity with spatial position and comparative anatomy, OTTER estimates correspondence weights across multiple candidate locations. The resulting mapping recovers established regional relationships, generates predicted human maps from mouse experimental responses and identifies candidate mouse targets from human maps of disease phenotype, symptoms and stage.

Mouse functional connectivity reconstructed human functional connectivity less accurately in expanded association cortex than in sensory and motor territories. This regional pattern was graded, with particularly low scores in the lateral prefrontal cortex. However, because the same connectomes contributed to fitting OTTER, reconstruction provides only an internal assessment of connectivity correspondence. In Control B, a separate analysis of 51 orthologous genes found a modest, non-significant reduction in best-match expression similarity relative to the other measured human parcels, with mouse matches selected independently of OTTER. The regional reconstruction pattern provides quantitative context for longstanding questions about correspondence between rodent and primate prefrontal cortex^3,4^.

Connectivity-based common spaces have previously enabled cortical maps to be translated between humans and macaques^11^. Our comparison with TransBrain^16^ illustrates how similar mean regional AUROCs can coexist with differences in prediction concentration and output sampling (Fig. 4). In the configurations evaluated here, OTTER generated parcel-wise predictions, while TransBrain returned one value per atlas region. OTTER’s correspondence weights also allow signals occupying different parts of the same source atlas region to produce distinct translated maps. These predictions can inform the selection of regions for injection, recording or stimulation, with spatial precision limited by the underlying measurements.

The disease applications show how anatomical priorities depend on the clinical feature being investigated. Alzheimer phenotype maps supported the prespecified mouse target sets in both discovery tau-PET and structural MRI analyses. The corresponding target assignments also showed selectivity in tau-defined subtypes in an external tau-PET cohort. Translation of the human TMS atlas^46^ yielded distinct candidate mouse regions for the dysphoric and anxiosomatic symptom domains examined in human stimulation studies^47^. In Parkinson disease, translated cross-sectional human stage maps showed an overall increase in the interoceptive-minus-motor contrast. The corresponding anatomical contrast showed an overall increase over time in a separate longitudinal mouse MRI analysis, motivating investigation of visceral, gustatory and agranular-insular regions alongside motor cortex. Together, these applications connect a specified clinical question to the choice of candidate anatomy for experiments.

The spatial detail of OTTER predictions is limited by the input connectomes and parcellation, and finer sampling cannot recover information absent from the underlying measurements. Regional benchmark AUROCs remained similar across cohort splits and refits omitting one connectivity modality, although the highest-weight human targets changed for individual mouse parcels. Fine-scale target selection therefore requires anatomical evidence and sensitivity analyses alongside the translated map. The alignment depends on curated anatomical anchors, which would need to be established for each additional species comparison. Its correspondence weights are conditional on these inputs and constraints and have not been calibrated as probabilities that individual correspondences are correct. The fixed reference mapping does not estimate disease- or participant-specific changes in correspondence. For applications based on published group-level maps, participant-level uncertainty in those source maps is not propagated through translation.

Targeted perturbation of the predicted mouse regions would test whether manipulating these regions changes specified behavioural, imaging or disease-related measurements in the chosen model. With suitable connectivity data and comparative anatomical guidance, OTTER could be extended to other species to generate corresponding regional predictions. OTTER translates mouse experimental maps into predicted human maps and identifies candidate mouse regions for testing hypotheses derived from human imaging.

## Methods

### Overview

To translate brain maps between mouse and human, we estimated a probabilistic coupling, π, that assigns weights to pairs of parcels across the two species. The coupling was fitted using semi-relaxed fused Gromov–Wasserstein optimal transport, combining within-species functional and structural connectivity with parcel coordinates and curated anatomical correspondences. We used the fitted coupling to translate maps in either direction by calculating weighted averages of measurements from source parcels. We evaluated anatomical correspondence using the full model and refits with selected anatomical constraints removed, and assessed translated biological maps against prespecified comparison maps.

### Human and mouse functional connectivity data

To estimate human functional connectivity, we used resting-state functional MRI (rs-fMRI) data from the Autism Brain Imaging Data Exchange (ABIDE) ^51^, accessed through the Preprocessed Connectomes Project^52^. The sample comprised 113 healthy controls aged 18–45 years (100 male; mean age, 28.1 years; SD, 5.6 years). We selected the C-PAC-preprocessed data in MNI152 space with 0.01–0.1 Hz band-pass filtering and global signal regression. Preprocessing also included slice-timing and motion correction, and nuisance regression incorporating motion parameters, CompCor components derived from white matter and cerebrospinal fluid, and linear and quadratic trends.

For mice, we used rs-fMRI data from the Grandjean resource^26^ and included 105 healthy C57BL/6 control animals (87 male; mean age, 6.3 ± 4.5 weeks). The repetition time was 1 s. Visual inspection revealed suboptimal anatomical registration in a subset of the downloaded images, so we reprocessed the dataset using RABIES^53^. This additional processing used the publicly available preprocessed images as inputs rather than raw MRI data. Functional images were nonlinearly registered to the DSURQE mouse brain template^54^ using symmetric normalisation (SyN) ^55^ and resampled to 0.2-mm isotropic voxels, with brain extraction and functional masking performed within RABIES. Data were band-pass filtered at 0.01–0.10 Hz, nuisance regressed using the six rigid-body motion parameters and aCompCor-derived regressors, and spatially smoothed with a 0.3-mm kernel.

#### Functional masks

To define the spatial coverage for region-of-interest (ROI) placement, we constructed a group functional mask for each species. Human binary masks were derived from the first volume of each preprocessed rs-fMRI dataset by retaining non-zero voxels and filling enclosed holes. Mouse binary masks were derived from each animal’s temporal standard-deviation map of the RABIES-preprocessed functional time series (func_bold_combined_STD_map.nii.gz), using Otsu’s thresholding^56^ followed by three-dimensional hole filling. Within each species, individual masks were averaged, and voxels included in at least 80% of masks were retained. The resulting group masks were in MNI152 space for humans and DSURQE space for mice.

#### Regions of interest

We selected 21 homologous anatomical classes from levels 3 and 4 of the multilevel atlas of Garin et al. ^23^ and represented each in both hemispheres, yielding 42 anchor ROIs per species. The final ROI sets contained 1,864 mouse ROIs and 2,094 human ROIs across both hemispheres, including these anchors. Equidistant ROI centres were placed within the respective group functional masks at approximately 0.5-mm spacing in mice and 10-mm spacing in humans. Centres were defined in physical coordinates (mm), and bilateral symmetry was enforced by mirroring them across the midsagittal plane. Spherical ROIs with radii of 0.35 mm in mice and 7 mm in humans were defined around these centres and restricted to voxels within the respective functional masks.

#### Functional connectivity

For each mouse and each human participant, we extracted the mean rs-fMRI time course within each ROI defined above. Pairwise Pearson correlations between these time courses yielded one ROI-by-ROI functional connectivity matrix per individual. Matrices were stacked separately for each species along the individual dimension, producing arrays of 1,864 × 1,864 × 105 for mice and 2,094 × 2,094 × 113 for humans (ROI × ROI × individual).

#### ROI lookup tables and atlas registration

We generated one ROI lookup table per species, shared across all individuals. Each table recorded ROI type, node ID, bilateral-pair ID, centre coordinates and voxel indices. Each ROI had a unique node ID, and bilateral partners shared a separate pair ID. Human centre coordinates were recorded in MNI152 space. The mouse table additionally included anatomical labels and atlas-based regional assignments.

To map mouse ROIs between atlas spaces, we nonlinearly registered the DSURQE template (0.07-mm isotropic voxels) to the Allen Mouse Brain Common Coordinate Framework template (CCFv3, 2017 release; 0.025-mm isotropic voxels) ^57^ and generated dense coordinate warp fields. For the equidistant mouse ROIs, anatomical labels were assigned from the DSURQE and Allen parcellations using both centre-based and majority-vote assignments, providing spatial reference and quality-control information.

### Mouse structural connectivity data

Mouse structural connectivity was derived from projection volumes aggregated within the summary structures of the Allen Mouse Connectivity Atlas^58^, accessed through the AllenSDK. We assigned each of the 1,864 mouse ROIs to an Allen summary structure using its centre voxel in CCFv3 space, following the atlas registration described above. The summary-structure connectivity matrix was symmetrised and represented on the ROI set using these assignments. ROIs assigned to the same summary structure therefore inherited identical structural connectivity profiles.

### Human structural connectivity data

Human structural connectivity was obtained from the group-averaged connectome of Domhof et al. 2022^59^ (HCP cohort, 10 million streamline count) defined on the Schaefer-400 17-network cortical parcellation^60^. The connectome was projected onto the 2,094-node parcellation by assigning each cortical node to the Schaefer parcel that occurred most frequently across its voxels. Thus, nodes assigned to the same Schaefer parcel inherited identical structural-connectivity fingerprints. Nodes outside the Schaefer cortical coverage were assigned zero structural connectivity in the projected matrix.

### Curated cross-species anatomical anchors

To guide cross-species alignment, we used two forms of anatomical constraint. The point anchors described above represented 21 anatomical classes selected from the atlas of Garin et al.^23^, with one anchor in each hemisphere per class, yielding 42 points per species. We also curated 26 regional correspondences from the comparative neuroanatomical literature, each linking mouse and human ROI sets judged to be homologous (Supplementary Table 1). Both types were incorporated as soft constraints in the cross-species feature cost, but only the point-anchor pairs were used to fit the spatial transformation from mouse to human.

### Fused Gromov–Wasserstein formulation

OTTER estimated a non-negative coupling π between n_m_ = 1,864 mouse ROIs and n_h_ = 2,094 human ROIs, where π_$%_ specifies the correspondence weight for mouse ROI (i) and human ROI (j). The coupling combined within-species connectivity relationships, encoded by relational cost matrices C^M^ and C^H^, with cross-species spatial and anatomical constraints, encoded by the feature cost matrix M. We estimated π by minimising the fused Gromov–Wasserstein objective^20^ subject to the marginal constraints described below:

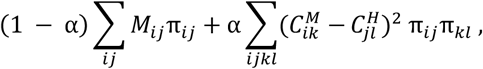

The feature term penalised cross-species pairings according to spatial distances and the point-anchor and regional constraints. The relational term compared pairs of mouse ROIs (i,k) with pairs of human ROIs (j,l), penalising discrepancies between *C^M^_ik_* and *C^M^_jl_* in proportion to π_ij_π*_kl_*. We set α = 0.5 for the full model, giving the two terms equal coeUicients.

### Within-species relational costs

To construct C^M^ and C^H^, we calculated functional and structural connectivity costs separately for each species. The functional cost C_FC_ was obtained from the group-mean ROI-by-ROI correlation matrix using (1-r), then symmetrised and assigned a zero diagonal. For structural connectivity, we applied log(1+x) to the connection weights x to compress their range. The transformed weights across the ROI set defined each ROI’s connectivity profile; correlations between these profiles were converted to the structural cost C_SC_ using (1-r).

Each cost matrix was divided by its largest oU-diagonal value, placing the functional and structural costs for both species in [0,1]. For the full model, we used fixed modality weights to define the relational cost for each species as C = 0.7 C_FC_ + 0.3 C_SC_.

### Cross-species feature cost

The cross-species feature cost M combined spatial distances with point-anchor and regional anatomical constraints. To calculate the spatial distances, we fitted a thin-plate-spline radial-basis transformation^65^ from the 42 mouse point anchors to their corresponding human coordinates (smoothing parameter 10^-3^). We calculated Euclidean distances between the transformed mouse ROI coordinates and the human ROI coordinates and normalised these distances to [0,1].

We initialised the cross-species cost matrix with the weighted, normalised spatial distances. Point-anchor constraints then replaced entries in the corresponding mouse rows and human columns with zero cost for the specified pairs and one for other pairings. Regional constraints next replaced entries whose mouse ROI or human ROI belonged to one of the 26 regional correspondences. Pairings whose mouse and human ROIs occurred together in at least one regional correspondence received zero cost; all other entries in these rows or columns received 0.15, while specified point-anchor pairs retained zero cost. Entries outside the supervised rows and columns retained their weighted spatial cost. The spatial weight was selected as described below; the point-anchor and regional penalties were fixed.

### Semirelaxed transport and solver

We used a semirelaxed formulation in which each mouse ROI assigned the same total correspondence weight, ∑_j_ *π_ij_* = 1/*n_m_*, while the total weight received by each human ROI was unconstrained. This allowed human ROIs to receive little transported mass without prescribing a human marginal distribution. We also examined a refit with the human marginal fixed and the mouse marginal unconstrained (Extended Data Fig. 9).

We fitted the coupling with the semirelaxed fused Gromov–Wasserstein solver in the POT library^64^, using the squared relational loss and mirror-descent updates. Each update multiplied the current coupling elementwise by the exponential of the negative objective gradient divided by *ε*, then rescaled its rows to restore the fixed mouse marginal. The parameter *ε* weighted a Kullback–Leibler penalty between successive iterates; the objective itself contained no entropy term. We initialised the coupling uniformly and used the 25th iterate at (*ε* =0.05).

### Regional benchmark and evaluation metrics

We evaluated regional correspondence using 19 mouse–human anatomical region pairs adapted from Beauchamp et al.^9^. Of 22 non-cerebellar candidate pairs, we retained those with a defined human mask and non-empty parcel sets in both species. Mouse parcels were assigned DSURQE labels using the most frequent non-zero label in a 5 × 5 × 5-voxel neighbourhood after translating their coordinates by (−0.027, −2.334, +1.018) mm; regions were formed from the corresponding DSURQE tree branches. Human target masks included parcels whose MNI centroids fell within spherical neighbourhoods of region-specific radius (6–15 mm), centred at curated left-hemisphere coordinates and their right-hemisphere reflections. This benchmark was used for model comparisons and parameter selection. Because it informed parameter selection and partly overlapped the anatomical supervision, we did not treat full-model performance as independent validation.

To evaluate a mouse–human region pair, we summed the coupling weights across mouse ROIs in the source region and ranked human ROIs by their received mass. We quantified regional correspondence using the area under the receiver operating characteristic curve (AUROC), which measured how well received mass separated ROIs in the expected human region from the remaining human ROIs. Mass-in-region was the fraction of the source region’s total transported mass received by the expected human region. For each mouse ROI in the source region, we separately ranked all human ROIs by coupling weight. Top-1, top-5 and top-10 recovery were the fractions of mouse ROIs with at least one expected-region human ROI among their first 1, 5 or 10 targets, respectively; mean rank averaged the rank of the highest-ranked expected-region human ROI across source mouse ROIs. Aggregate AUROC, mass-in-region and top-k recovery were weighted across benchmark pairs by the number of source mouse ROIs.

Regional localisation error was the Euclidean distance, in millimetres, between the mass-weighted centroid of the received human distribution and the centroid of the expected human region. We assessed preservation of spatial relationships by correlating pairwise distances between mouse ROIs with the distances between their corresponding mass-weighted human centroids.

For each benchmark pair, we evaluated AUROC, mass-in-region, top-1 recovery and centroid displacement using the metrics defined above. We tested transported-mass enrichment against 1,000 samples of human parcels drawn uniformly with replacement, each with as many entries as the expected region. Centroid displacement was tested against 1,000 uniformly random three-dimensional rotations of the expected-region centroid about the mean of all human parcel centroids, keeping the predicted centroid fixed. One-sided Monte Carlo P values counted sampled mass values at least as large as observed or null displacements no greater than observed, with a plus-one correction. P values were adjusted using the Benjamini–Hochberg procedure separately across the 19 regions for each test (false discovery rate, 5%).

### Solver regularisation and parameter selection

We selected the solver parameter *ε* and the spatial-cost weight using the 19 mouse–human region pairs in the regional benchmark described above. For each pair, top-1 accuracy was the fraction of source mouse ROIs whose highest-weight human target lay in the corresponding human region. Parameter combinations were compared using the unweighted mean of these per-pair scores. All other model settings were fixed.

We evaluated 25 parameter combinations, with *ε* ∈ 0.005,0.02,0.05,0.1,0.2 and spatial weights ∈ 0.1,0.25,0.5,0.75,1.0. The 19 benchmark pairs were divided into five folds for scoring. For each fold, we selected the combination with the highest mean per-pair top-1 accuracy on the remaining pairs and evaluated it on the held-out pairs. The mean of the five held-out fold scores was 0.673. This score evaluated the accuracy-based parameter-selection procedure; it did not constitute independent biological validation because some benchmark regions overlapped the curated anatomical constraints used to fit OTTER.

For the full model, we used a spatial weight of 0.25 and *ε* =0.05. At this spatial weight, *ε* =0.05 and *ε* =0.2 gave similar mean per-pair top-1 accuracy (0.670 and 0.674, respectively). We selected *ε* =0.05 because its correspondence weights were more concentrated than at *ε* =0.2.

### Cost-term ablation and supervision withholding

To assess how the cost components affected regional correspondence, we refitted OTTER with four combinations of costs. Starting with the relational cost alone, we successively added the spatial cost derived from the point-anchor transformation, direct point-anchor costs and regional correspondence costs. We evaluated all four variants on the 19-pair regional benchmark using the metrics defined above (Fig. 2a).

To assess recovery when supervision was withheld, we refitted the full model after omitting each of 41 supervision units in turn: 15 scorable Garin classes and 26 regional correspondences. Each refit retained the connectivity and spatial costs and the remaining anatomical supervision, and was evaluated on the omitted unit (Fig. 2d). For each omitted unit, we removed all mouse and human point anchors sharing a pair identifier with a mouse anchor in that unit from the direct anchor cost and the thin-plate-spline fit. We refitted the transformation from the remaining pairs and removed regional correspondences for which more than half of their mouse parcels lay within the omitted unit.

### Target-wise anatomical supervision withholding

To assess recovery after removing anatomical supervision for a target, we withheld all point-anchor and regional supervision overlapping each benchmark region in turn and refitted OTTER. Where the omitted point anchors had contributed to the spatial transformation, we refitted that transformation from the remaining pairs. With the selected hyperparameters fixed, we compared three configurations: connectivity and spatial costs together, spatial cost without connectivity, and connectivity without the spatial cost. Each configuration retained anatomical supervision outside the withheld region (Fig. 2c; Extended Data Fig. 4).

### Cross-species translation of scalar maps

Maps containing one numerical value per parcel were translated between species using coupling-weighted averages. For mouse-to-human translation, the weights were normalised separately for each human parcel; for human-to-mouse translation, they were normalised separately for each mouse parcel. Source-map sampling and missing-data treatment are described for each analysis below.

### Biological map comparisons

To assess cross-species correspondence, we evaluated fourteen biological maps: one myelin-associated functional-connectivity gradient, the T1w/T2w ratio and cytoarchitectural type, eight cell-class marker-expression maps and contrasts, and three layer-marker expression contrasts^27–29,66,67^.

For the functional-connectivity gradient, T1w/T2w ratio and cytoarchitectural-type comparisons, mouse parcels without measurements were excluded from the weighted average, and the weights were renormalised over the remaining mouse parcels for each human target. Each translated map was compared with its prespecified human comparison map using Pearson correlation over human parcels with finite translated and comparison values. The primary gene-expression comparisons were restricted to human parcels containing donor tissue. Correlations were tested against the translation-spin null described below.

#### Functional-connectivity gradients

We derived the first four diffusion-map components separately from the mouse and human functional connectomes. Within each species, we selected the component with the largest absolute Spearman correlation with its T1w/T2w ratio map; the second component was selected in both species. We translated the selected mouse gradient and compared it with the selected human gradient using the absolute Pearson correlation.

This comparison was an internal test of connectivity preservation because the gradients were derived from the functional connectomes used to fit OTTER. We repeated the selected-gradient comparison after refitting OTTER without the connectivity cost (α = 0), and after randomly reassigning mouse parcels to rows of the fitted coupling. We also translated all four mouse components with the full coupling and the coupling fitted without connectivity, calculating their absolute Pearson correlations with each of the four human components (Extended Data Fig. 6).

#### Marker-expression maps

Cell-class marker-expression maps were constructed from a curated panel of 61 marker genes, with class assignments based on published cell-type atlases^67–69^. Mouse expression was sampled from Allen in situ hybridisation energy volumes at 200-µm resolution^61^, and human expression was obtained from the Allen Human Brain Atlas^63^ and processed with abagen^62^. These maps represented regional marker-gene expression rather than direct estimates of cell abundance.

Expression measurements were available for 1,713 of 1,864 mouse parcels and 1,040 of 2,094 human parcels. The primary cell-class comparisons used the 1,040 human parcels containing donor tissue. Of these, 860 were cortical, out of 1,768 cortical parcels in the human parcellation. The layer-marker comparisons used this cortical subset and evaluated three contrasts: granular layer 4 minus infragranular, supragranular minus infragranular, and supragranular minus granular marker expression.

For all eleven marker-expression measures, missing expression values were replaced by each gene’s mean across measured parcels within the same species before gene-wise z-scoring and map construction. The human donor-tissue mask was defined before imputation. We also evaluated the eight cell-class measures over all 2,094 human parcels and the three layer contrasts over all 1,768 human parcels assigned to the Schaefer-400 cortical atlas, including those without donor tissue. A cortex-only sensitivity analysis evaluated glutamatergic expression, the excitatory–inhibitory and neuronal–glial contrasts, and the dopaminergic marker map. Human scoring was restricted to 856 donor-tissue parcels with centres within 6 mm of a labelled Harvard–Oxford cortical voxel, with all 1,864 mouse parcels retained for translation.

### Network-level correspondence

Mouse parcels were assigned to eleven resting-state networks using curated labels for the Garin anatomical classes, propagated to other parcels from the nearest anchor. Ten networks with specified human counterparts were evaluated against human reference labels based on the Yeo cortical parcellation^31^ and a separate subcortical category. Because the mouse labels were derived from anatomical classes used to guide OTTER, this analysis assessed network-level consistency of the fitted correspondence.

For each mouse network, coupling weights were summed over its source parcels to obtain its distribution of transported mass across human parcels. We summed this mass across human parcels within each reference network and counted how many of the ten expected counterparts ranked highest. This count was tested against 500 rotations of the discrete mouse network labels, with translation and scoring repeated through the fixed coupling (one-sided source-label spin test). For the spatial display, each human parcel was assigned to the mouse network contributing the greatest total coupling mass to that parcel (Fig. 3b). For the compactness analysis, we selected each mouse parcel’s highest-weight human target parcel and calculated, within each of the eleven mouse networks, the mean Euclidean distance of these targets from their centroid (mm). Each observed spread was compared with the mean spread from 100 coupling-row permutations.

We derived seven mouse network maps by applying FastICA (seed 42; maximum 2,000 iterations) to the same functional-connectivity matrix used to fit OTTER. Correlations were Fisher z-transformed, the diagonal was set to zero and values below each row’s 90th percentile were set to zero. Component signs were oriented so that the loading with the largest absolute magnitude was positive. Each component was labelled by the most frequent curated mouse network among parcels in its highest 5% of weights. We multiplied each signed component map by the fixed coupling matrix, ranked human reference networks by the mean resulting weight across their parcels and counted a match when the specified counterpart ranked highest. Six components had specified human counterparts; the brainstem-labelled component was excluded from the match count.

### Spatial null models and permutation controls

To assess spatial correspondence between brain maps, we generated rotation-based null maps using an adaptation of cortical spin methods^17,70^. In the default procedure, parcel centroids were projected onto a sphere, and each null draw applied the same uniformly random three-dimensional rotation to all centroids. Map values were reassigned by matching each original parcel independently to its nearest rotated parcel.

For cortical-map spin tests, one map in the cortical comparison was rotated while the other remained fixed. Target-map spin tests of mouse-to-human correspondence rotated the human comparison map while keeping the translated mouse map fixed. Translation-spin tests rotated the source map before translation through the fixed coupling, then compared the translated map with the unchanged target map. This test evaluated correspondence conditional on the fitted coupling, whose spatial structure could produce a non-zero null correlation.

Unless otherwise specified, rotation-based brain-map correlation tests used N = 1,000 rotations and two-sided Monte Carlo P values calculated as P = (B + 1)/(N + 1), where B was the number of rotations with an absolute correlation at least as large as observed. The tests in Extended Data Fig. 7 used whole-sphere rotations; analysis-specific rotation constraints and alternative spatial surrogates are described in the relevant Methods sections.

Non-spatial controls randomised other parts of the analysis. For the benchmark mass null, human parcels were sampled uniformly with replacement, matching the number of parcels in the expected region. The anchor-pair permutation null shuffled anatomical correspondences before refitting the coupling. The coupling-row permutation null reassigned mouse parcels to rows of the fixed coupling, whereas the source-region-value permutation null shuffled measurements among named source regions before retranslation. These controls did not impose spatial-autocorrelation constraints.

The processed mouse input maps contained 1,864 distinct values for the myelin-associated functional-connectivity gradient and 39 for the T1w/T2w ratio map; the cytoarchitectural-type map encoded five ordinal categories^28^. After translation, these maps were compared with their human comparison maps over 2,094, 1,789 and 1,787 human parcels, respectively (Extended Data Fig. 7). These parcel counts described the number of locations included in each correlation, without implying statistical independence. For these three comparisons, we reported P-values from the spatial null models described above; parametric correlation P values were not reported.

### Reconstruction accuracy

To quantify how well mouse functional connectivity reproduced human connectivity, we reconstructed the human functional-connectivity matrix through the fitted coupling. For human parcels with non-zero received weight, we defined a human-by-mouse weight matrix A, with *A*_ji_ = *π*_ij_/ ∑_k_ *π*_ij_, so that each row summed to one. The reconstructed matrix was

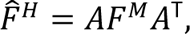

where *F^M^*^’^ was the group-mean mouse functional-connectivity matrix. Each human parcel’s reconstruction score was the Pearson correlation between its reconstructed and measured group-mean human connectivity profiles across all other human parcels, using finite values in both profiles and excluding the diagonal. Undefined correlations were excluded. As these connectomes also contributed to fitting OTTER, reconstruction provided an internal assessment of connectivity correspondence.

We assessed hemispheric consistency using Spearman correlations between reconstruction scores in matched left and right parcels and between mean scores in matched Schaefer areas. We also repeated reconstruction using both species’ structural-connectivity matrices while keeping the fitted coupling fixed (Extended Data Fig. 5a).

Human measurement reliability was assessed by splitting the cohort into two halves and correlating each parcel’s functional-connectivity profiles estimated separately in each half. We then divided each parcel’s reconstruction score by its split-half reliability to obtain a normalised score (Extended Data Fig. 5b).

To assess reconstruction against spatially rearranged mouse connectivity, we generated 200 surrogate mouse connectomes. For each surrogate, mouse parcel coordinates were randomly rotated, and each rotated parcel was assigned its nearest original parcel. The resulting index map was applied to both rows and columns of the mouse functional-connectivity matrix. Each surrogate was reconstructed through the unchanged coupling, with human functional connectivity held fixed. We compared observed mean reconstruction accuracy, the expansion correlation and the Control B contrast defined below with their respective surrogate distributions (Extended Data Fig. 5c).

To characterise the spatial distribution of reconstruction accuracy, we averaged parcel scores within each represented Schaefer-400 region and calculated Spearman correlations with seven reference maps. These comprised macaque-to-human cortical expansion estimated by Hill et al.^32^ and Xu et al. ^14^, human postnatal cortical expansion^32^, the macaque–human functional-connectivity homology index^14^, the sensorimotor–association axis^30^, the principal functional-connectivity gradient^27^ and the T1w/T2w ratio^29^. Each comparison included regions with values available for both measures and was assessed using the cortical-map spin null.

To compare reconstruction at opposite ends of the sensorimotor–association axis, we ranked Schaefer regions by their axis values and selected equal-sized lowest and highest thirds. These groups represented the sensorimotor and association tertiles, respectively. We reported the sensorimotor-minus-association difference in mean reconstruction accuracy and Cohen’s d, and assessed the mean difference using cortical-map spin rotations of the regional reconstruction scores.

We also summarised parcel scores within the 16 network labels represented in the supplied Schaefer-400 table, based on the Yeo 17-network parcellation^31^. This table combined the two limbic subdivisions under one Limbic label. We included 1,824 cortical parcels with finite reconstruction scores and T1w/T2w values. For the network map, we subtracted the overall parcel mean from each network’s mean reconstruction score and divided by the population standard deviation across these parcels. Each network contrast was the difference between its mean score and the mean across all other included parcels, divided by the same standard deviation. Contrasts were assessed using 2,000 cortical-map spin rotations (two-sided), with Bonferroni correction across the 16 networks containing at least 30 parcels.

### Transcriptomic comparison

To compare transcriptomic similarity with connectivity reconstruction, we assembled expression profiles for 51 one-to-one orthologous genes in mouse and human. Mouse expression was obtained from Allen in situ hybridisation energy volumes at 200-µm resolution^61^ and sampled at each ROI using the DSURQE-to-CCFv3 transformation described above. Human expression was obtained from the Allen Human Brain Atlas^63^ and processed with abagen^62^. These data were represented as mouse and human ROI-by-gene matrices of 1,864 × 51 and 2,094 × 51, respectively. Note that these expression data were not used to fit the OTTER coupling.

To compare connectivity reconstruction with transcriptomic similarity, we z-standardised each gene using all available parcel measurements, separately within each species. Parcels with missing expression measurements for any of the 51 genes were excluded without imputation, leaving 1,763 eligible mouse parcels. Each parcel’s gene profile was then centred across genes and normalised to unit length. For each eligible human parcel, we calculated cosine similarity with all eligible mouse profiles and retained the maximum value. This mouse match was selected from gene expression without using the OTTER coupling. For the Control B analysis, both measures were evaluated on the same 1,040 human parcels with complete expression profiles and finite reconstruction scores. Each contrast was the mean in the 43 Control B parcels minus the mean across the other 997 parcels, divided by that measure’s population standard deviation across all 1,040 parcels. Contrasts were assessed separately using the same 2,000 spatial spin rotations on this shared parcel set (two-sided).

### Comparison with TransBrain

To compare regional correspondence on a common anatomical scale, we evaluated OTTER and the TransBrain Python package^16^ over 127 shared human regions from the Brainnetome-based configuration^71^. The 24 mouse queries were selected from the reference correspondence tables distributed with TransBrain’s code, retaining queries covered by both methods with at least one reference target among the common human regions. For each mouse query, we summed OTTER weights over source parcels and then over human parcels within each target region. TransBrain queries assigned a value of one to the source mouse region and zero to the other regions. Negative TransBrain scores were set to zero, and vectors with positive total weight were normalised to sum to one over the common human region set. OTTER was not fitted to TransBrain outputs or transcriptomic embeddings. However, some queried mouse regions overlapped OTTER’s regional correspondence constraints on the mouse side, so this benchmark was not fully held out from anatomical supervision.

For each of the 24 mouse-region queries, we calculated AUROC, top-1, top-3 and top-5 recovery, and the fraction of normalised weight assigned to the reference human regions. AUROC quantified how well scores separated reference regions from the remaining targets. Top-k recovery was successful if at least one reference region was among the k highest-scoring targets. Metrics were averaged across queries, and per-query AUROC values were compared using a paired Wilcoxon signed-rank test. Prediction concentration was quantified as *N = 1/Σ_h_p^2^_h_*, where *p_h_* was the normalised weight of human target ℎ on the same 127-region set. Effective counts were averaged across queries with non-zero total weight.

Translation self-consistency was assessed by translating three regional mouse maps into human space and then back into mouse space: the functional-connectivity gradient averaged within mouse regions, anterior-insula optogenetic activation and the Magel2 map. For OTTER, available regional input values were assigned to parcels with the corresponding region label, and translation used the finite-source weighted averages described above. TransBrain received a complete mouse-region input vector, with missing values replaced by the mean available value across its mouse regions for that map; imputed entries were excluded from scoring. We averaged OTTER’s returned parcel values within mouse regions and calculated Pearson correlations with the original regional values. Both methods were scored on the same regions with measured input values and OTTER parcel coverage, giving the same 52 of the 68 configured mouse regions for all three maps.

For the error maps in Fig. 4b, original and returned maps were separately z-scored on their common finite parcel mask. Errors were returned minus original values, and root-mean-square error was calculated on that mask.

Forward-gradient correspondence was assessed using absolute Pearson correlations between each translated map and the observed human gradient, and between the two translated maps, over Brainnetome cortical regions represented in all three maps.

### Translation of a mouse optogenetic activation map

To assess mouse-to-human translation of an experimental activation map, we translated the response to optogenetic stimulation of the mouse agranular insula^33^ using the forward procedure described above (Fig. 6a). Regional input values were assigned to all mouse parcels with the corresponding Allen structure acronym; parcels without finite input values were excluded from the forward weighted average. The human prediction was z-standardised using the mean and population standard deviation across its 2,059 finite parcels, then summarised by mean values within the 16 represented Yeo-17 network labels (Fig. 6b). Enrichment was the mean z-score across the 261 parcels labelled Salience/Ventral Attention A or B minus the mean across the remaining 1,798 finite parcels, including those without a network label; each parcel was weighted equally. For the OTTER-only analysis, we assessed this enrichment using 1,000 coupling-row permutations, which randomised the assignment of mouse parcels to rows of the fixed coupling matrix. Each permuted map was z-standardised across its finite predictions, and the contrast was recalculated on the original parcel sets using finite permuted values and assessed with a one-sided upper-tail test.

For comparison with TransBrain, we evaluated both translated maps on the same 1,635 human parcels (Fig. 6c). Each human parcel’s Brainnetome label was sampled at the atlas voxel nearest its coordinates; background labels were replaced by the modal nonzero label in the surrounding 3 × 3 × 3 voxels, when available. TransBrain regional predictions were repeated at the human parcels assigned to each Brainnetome region. The shared set required both a Yeo-17 label and a Brainnetome label, with finite predictions from both methods. Each method’s prediction was z-standardised separately on this set using its mean and population standard deviation. We calculated the mean z-score across Salience/Ventral Attention A and B parcels minus the mean across the other included parcels, weighting parcels equally. Each method was then evaluated using 1,000 source-region-value permutations: activation values were shuUled among the named mouse input regions, the permuted map was translated again, and each prediction was standardised again and enrichment was recalculated on the fixed common human parcel set. These tests assessed each method’s enrichment relative to its own permuted-input distribution (one-sided upper-tail tests).

### Reverse translation of functional and molecular maps

Human volumetric maps were resampled to the grid of the 2,094-parcel human parcellation using linear interpolation. We averaged map values within each parcel and assigned the arithmetic mean of the finite parcel means to parcels without finite values. These parcel values were translated to the mouse using the fitted coupling and the reverse weighted average defined above. The translated values were then averaged within each mouse anatomical structure.

#### Functional-system maps

To evaluate reverse translation of functional maps, we used human meta-analytic activation maps for twelve functions and behaviours (Supplementary Table 2). For each system, we generated a map from Neurosynth v7 using NiMARE’s multilevel kernel density analysis chi-square estimator^34,73,74^. Studies were selected using the corresponding Neurosynth term with a term frequency–inverse document frequency (tf–idf) threshold of 0.001. We used the estimator’s specificity (association-test) output. Volumetric maps in MNI152 space were used to include subcortical targets, which are not represented in cortical surface maps. The systems, their Neurosynth terms and expected mouse structures were fixed before scoring; the expected structures and supporting references are listed in Supplementary Table 2.

For functional-system ranking, we retained structures represented by at least five parcels, leaving 83 of the 166 structures in the mouse parcellation, and ranked them by their mean translated value. The test statistic for each system was the highest score among its prespecified expected mouse structures. We compared this statistic with a reverse translation-spin null generated by rotating the human source map, reassigning values using the nearest-centroid procedure described above, and repeating translation through the fixed coupling. Each of the 1,000 rotations was scored using the same expected structures, and significance was assessed by counting null statistics that met or exceeded the observed statistic. Recovery was defined as at least one eligible expected target ranking among the top three of these 83 structures.

#### Molecular maps

To evaluate reverse translation of molecular maps, we selected volumetric MNI152 annotations from neuromaps by tracer name^36,37^ and applied the preparation and translation procedure described above. The eight dopamine maps covered D1 receptor binding (SCH23390), D2/3 receptor binding (fallypride, FLB457 in two datasets and raclopride in two datasets), and dopamine transporters (FE-PE2I and FP-CIT). Reference maps covered serotonin (nine maps), μ-opioid receptors (two maps), cannabinoid CB1 receptors (one map) and GABA-A receptors (two maps).

For each translated map, we calculated the mean predicted value within each of the 83 mouse structures represented by at least five parcels and replaced negative structure means with zero. Striatal mass fraction was the sum of these non-negative structure means in caudoputamen, nucleus accumbens and olfactory tubercle divided by the sum across all 83 structures, giving each structure equal weight; a zero denominator gave a fraction of zero. Separately, for each dopamine map, we tested the maximum of the unclipped mean predictions in caudoputamen, nucleus accumbens and substantia nigra pars reticulata against the same statistic after 1,000 rotations of the human input map and reverse translation through the fixed coupling (one-sided upper-tail test). We tested whether dopamine-map fractions were higher than those of the pooled reference maps using an exact one-sided Mann–Whitney U test (Fig. 6f).

### Alzheimer phenotype reverse translation

To compare translation across Alzheimer phenotypes, we used tau-PET and voxel-based morphometry T maps from NeuroVault collection 8546^40^. The tau-PET maps (image IDs 402706–402708) and morphometry maps (402719–402721) corresponded to AD, PCA and lvPPA, in that order. Positive source T values indicated greater tau binding or lower grey-matter volume in the named phenotype; we applied no sign reversal or additional thresholding. These maps were derived from 119 amyloid-positive participants with PCA, lvPPA or AD without PCA or lvPPA, with each group contrasted against the other two. The original tau-PET models adjusted for age, Clinical Dementia Rating Sum of Boxes and global cortical tau-PET standardised uptake value ratio (SUVR); the morphometry models adjusted for age, Clinical Dementia Rating Sum of Boxes and global cortical grey-matter volume.

Human OTTER parcels were assigned to Desikan–Killiany cortical regions across both hemispheres^75^. Atlas labels were sampled at the voxel nearest each parcel’s coordinates. For background labels, we used the nearest cortical atlas voxel within four atlas voxel units; parcels with non-cortical labels or no cortical assignment were excluded. We retained 52 regions represented by at least eight parcels, keeping hemispheres separate. For each map, we averaged parcel values within each retained region and z-standardised the 52 regional means using their mean and sample standard deviation. The human parcel weights for each mouse target set (defined below) were summed within each retained region, and the target score was the weighted sum of regional z-scores. We did not renormalise the weights over the retained regions.

Before scoring, we specified three mouse target sets: entorhinal cortex and hippocampus for AD without PCA or lvPPA (ENTl, ENTm, CA1, CA3, DG and SUB); visual cortex for PCA (VISp, VISl, VISal, VISam, VISpm, VISrl, VISa and VISpor); and auditory and temporal association cortex for lvPPA (AUDp, AUDd, AUDv and TEa). The lvPPA assignment was evaluated as an auditory–temporal target hypothesis. Mouse TEa includes auditory circuitry that receives input from primary auditory cortex and participates in processing vocalisations.

For the primary analysis, we averaged row-normalised coupling weights across all mouse parcels within each target set, giving each parcel equal weight. These mean weights were used to score each human phenotype map for each target set. In a sensitivity analysis, we first averaged weights within each anatomical structure and then across structures, giving each structure equal weight. For each phenotype map, selectivity was the score for its assigned target set minus the mean of its scores for the other two sets. Joint selectivity (C) was the sum of the three phenotype selectivities.

We assessed joint selectivity separately for tau-PET and voxel-based morphometry using a one-sided test for C values greater than expected under a null distribution generated from 10,000 spatial rotations. Rotations were mirrored across hemispheres, with one-to-one reassignment within each hemisphere; the same reassignment was applied to all three phenotype maps before translation through the fixed coupling and recalculation of C. We repeated the analysis after refitting OTTER without the eight directly relevant entorhinal/hippocampal, visual and auditory regional correspondence packs, retaining the remaining anatomical supervision and spatial warp.

#### LEADS tau-subtype analysis

To evaluate the same target assignments in an external dataset, we used baseline flortaucipir PET maps from the Longitudinal Early-Onset Alzheimer’s Disease Study (LEADS; NeuroVault collection 23001) ^45^. The primary inputs were the authors’ subtype-versus-rest T maps for the typical (S1), posterior (S3) and left-temporal (S2) tau subtypes, identified using Subtype and Stage Inference (SuStaIn) ^76^. These maps were adjusted for SuStaIn stage, age, sex, education and amyloid burden (Centiloid), and thresholded at uncorrected P < 0.001. The assigned mouse targets were entorhinal cortex and hippocampus for S1, visual cortex for S3, and auditory and temporal association cortex for S2. Before inspecting the results, we defined confirmation as positive selectivity for all three subtypes and joint one-sided P < 0.05 under both full OTTER and the refit without the eight relevant regional correspondence packs.

For the unthresholded sensitivity analysis, we used subtype mean standardised uptake value ratio (SUVR) maps within SuStaIn stage bins 9–11, 12–14 and 15–20. Within each bin, we subtracted the equally weighted mean of the other two subtypes’ maps from each subtype’s map. These contrast maps were z-standardised and then averaged across the three bins. We applied the same selectivity statistic and spatial-null test under both couplings. The stage 0–8 bin was excluded because it contained six participants in the left-temporal subtype.

### TMS symptom-circuit dissociation

We obtained the human dysphoric–anxiosomatic TMS atlas from NeuroVault collection 13075 (image 787858)^77^. Negative values defined the dysphoric input and positive values the anxiosomatic input. To account for non-finite source voxels, interpolation weights were renormalised over voxels with finite values. We averaged the sampled values within 2,086 human parcels; the remaining eight parcels, which were absent from the voxel parcellation, were sampled at their MNI centroids using the same normalisation. For each sign, we constructed a non-negative input map and defined its upper 5%, 10%, 20% and 30% tails using percentiles across all 2,094 human parcels, including values set to zero for the opposite sign. We retained the amount by which each value exceeded the corresponding percentile threshold and set values at or below it to zero. The 10% threshold was used for the primary analysis.

We defined a medial prefrontal and cingulate target set for the dysphoric input (PL, ILA, ACAd, ACAv, ORBm and FRP), and an amygdala and insula target set for the anxiosomatic input (BLA, BMA, CEA, LA, AId, AIv and AIp). All listed structures were retained, including ILA and LA. Each translated input map was scored against both target sets. For the primary score, we summed translated values across the mouse parcels in each target set and divided by the sum across all named mouse parcels. For the structure-balanced sensitivity analysis, we first averaged translated values within each anatomical structure, then divided the sum of these means within a target set by their sum across all named structures. For each input, bias was the medial prefrontal/cingulate score minus the amygdala/insula score. The contrast between inputs C was the dysphoric bias minus the anxiosomatic bias.

We generated 10,000 surrogate human maps using joint bilateral-pair Moran spectral randomisation (seed 20260826). Each bilateral pair was represented by the mean of its left centroid and its right centroid reflected across the midsagittal plane. Left and right map values were treated as two features, with the same random sign applied to both features for each Moran mode. Separate Moran bases were fitted to cortical and non-cortical pairs.

The same surrogate maps were used for full OTTER, the refit without the amygdala, cingulate and lateral-prefrontal regional correspondence packs, and the matched coupling without the connectivity cost. One-sided tests evaluated C for each coupling and the paired difference in C between full OTTER and the coupling without connectivity (full OTTER minus no connectivity). For each comparison, maximum-statistic correction controlled the family-wise error rate across the 5%, 10%, 20% and 30% tail thresholds. Monte Carlo P values were calculated as (B + 1)/(10,000 + 1), where B counted null statistics at least as large as observed, including numerical ties. For threshold-corrected tests, these counts used the corresponding maximum-statistic null distribution.

For the threshold-free analysis, we retained the full non-negative input map for each sign without subtracting a percentile threshold and used the whole-brain normalisation defined above. Each input’s target-conditional bias used the same target-difference numerator but replaced the whole-brain denominator with the total within the two target sets, using the corresponding parcel or structure weighting. We evaluated these conditional biases for both the primary 10% tails and threshold-free inputs. All thresholded, threshold-free and target-conditional analyses were repeated with structure-balanced weighting for each of the three couplings and the paired full-minus-no-connectivity comparison. These analyses used the same surrogate maps and one-sided tests; maximum-statistic correction remained confined to the four tail thresholds.

### Parkinson disease stage translation

We obtained regional cortical-thickness effect maps for Hoehn–Yahr stages 1, 2, 3 and 4/5 from the cross-sectional ENIGMA-Parkinson study^48^. Each stage group was compared with age- and sex-matched controls. We negated the reported effect sizes *d* so that positive values denoted lower cortical thickness relative to controls. These values were mapped to 52 Desikan–Killiany regions across both hemispheres and z-standardised within each stage map.

We formed a linear stage map by weighting the four standardised stage maps by centred coefficients −1.5, −0.5, 0.5 and 1.5, respectively, and dividing the weighted sum by five, the sum of squared coefficients. Each stage map used the sample standard deviation across the 52 human regions; the four ordered stage groups were treated as equally spaced. Using OTTER, we compared translated scores for visceral, gustatory and agranular-insular regions (VISC, GU, AId, AIv and AIp; the interoceptive set) with those for primary and secondary motor cortex (MOp and MOs), giving equal weight to structures within each set. Coupling weights were row-normalised over all human parcels, averaged within mouse structures and equally across structures in each target set, then summed within the retained human regions. We multiplied each region’s linear-stage value by its interoceptive-minus-motor weight and summed across regions, without renormalising weights over the retained regions.

Significance of the trend under full OTTER was assessed using 10,000 spatial rotations, mirrored across hemispheres with one-to-one reassignment within each hemisphere, and a two-sided test. Linear trends were also assessed for each hemisphere, the direct human insula-minus-precentral contrast and each alternative coupling (10,000 rotations; two-sided tests). Hemisphere analyses re-standardised each stage map within that hemisphere and used separate one-to-one rotations; the bilateral analyses shared the primary rotations. Paired one-sided tests assessed whether full OTTER produced greater linear trends than the connectivity-only coupling, the spatial-only coupling and the coupling with spatial and anatomical costs but no connectivity cost, using full-minus-comparator trend differences under the shared bilateral rotations. Individual stage scores were descriptive.

### Parkinson disease stage analysis in QPN-NC

We assessed the stage-related pattern using the restricted Quebec Parkinson Network Neuroimaging Cohort (QPN-NC), a cross-sectional cohort of 202 participants with Parkinson disease and 69 older controls. Using the released demographic, diagnostic, clinical and FreeSurfer-derived structural data, we included all 69 controls and 141 participants with Parkinson disease who had cortical-thickness measurements and a Hoehn–Yahr assessment. For participants with multiple clinical assessments, we selected the Hoehn–Yahr assessment closest to MRI among those with available stage and interval data; all selected assessments were within six months of MRI. The signed clinical–MRI interval, measured in months and positive when the clinical assessment followed MRI, was included as a covariate in the stage-association analyses. For the ordinal analyses, Hoehn–Yahr stages 4 and 5 were pooled to give four ordered levels (1, 2, 3 and 4/5).

For each of the 50 cortical regions shared by QPN-NC and the ENIGMA analysis, we modelled cortical thickness in controls as a function of MRI age and sex. We used these models to calculate individual thinning maps for participants with Parkinson disease as predicted minus observed thickness, divided by the control residual standard deviation for each region. Positive values therefore indicated lower thickness than predicted from the controls.

At each region, we estimated the association between thinning and four-level Hoehn–Yahr stage across participants with Parkinson disease using ordinary least-squares regression, adjusting for MRI age, sex and clinical–MRI assessment lag. We z-standardised the resulting stage coefficients across the 50 regions using their sample standard deviation. We restricted the regional contrast weights defined above to these 50 regions without renormalising them. The contrast D was the sum of each region’s standardised coefficient multiplied by its interoceptive-minus-motor weight. A one-sided spatial test compared the observed D with values obtained after 10,000 hemisphere-preserving, one-to-one rotations of the human stage-coefficient map.

Each participant’s thinning map was z-standardised across the 50 cortical regions and scored using the same fixed OTTER contrast, yielding one score per participant. We assessed the association of these scores with four-level Hoehn–Yahr stage using partial Spearman correlation, adjusting ranked scores and stage for ranked covariates (MRI age, sex indicators and clinical–MRI assessment lag). For each of 10,000 mirrored spatial rotations, we reassigned regional values one-to-one within each hemisphere, applying the same reassignment to every participant’s map while keeping the contrast weights, stage and covariates fixed. We recalculated participant scores and their partial Spearman correlation with stage for each rotation and used a one-sided upper-tail test.

We examined three alternatives to the primary four-level regional model: ordinary least-squares regression using raw stages 1–5 or stages 3–5 versus 1–2, and Huber regression using four-level stage. An omnibus test combined these four regional analyses with the participant-level association using synchronised spatial rotations. For each of the five statistics, we centred the observed and rotated values on that statistic’s rotation-null mean and divided by its rotation-null sample standard deviation. We compared the largest standardised observed statistic with the largest standardised statistic in each rotation using a one-sided upper-tail test.

We assessed influence on the regional contrast D by refitting the regional analysis after excluding each participant with Parkinson disease or each control in turn. For control exclusions, we first refitted the control reference models and recalculated individual thinning maps.

#### Destrieux parcellation sensitivity analysis

To assess sensitivity to cortical parcellation, we used Destrieux cortical-thickness measures from the FreeSurfer v7.3.2 outputs. Mean thickness was extracted from aparc.a2009s.stats for 74 cortical regions per hemisphere, excluding the medial wall (148 regions in total). Reconstructions were excluded if recon-all failed or any Destrieux region was missing; missing thickness values were not imputed. The single excluded reconstruction was outside the primary sample, which remained 69 controls and 141 participants with Parkinson disease.

We represented the fixed OTTER contrast in this parcellation by assigning human cortical nodes to the nearest same-hemisphere fsaverage5 pial vertex within 8 mm. The contrast weights of the 1,580 mapped nodes were summed within their assigned Destrieux regions. Nodes not assigned a valid Destrieux region were excluded, and retained weights were not renormalised.

We repeated the regional and participant-level analyses, along with the regional Huber regression and leave-one-out checks. One-sided spatial inference used 10,000 hemisphere-preserving, one-to-one rotations of the Destrieux parcel centroids. Sampling uncertainty was estimated using 10,000 bootstrap resamples, with controls and participants with Parkinson disease drawn independently with replacement. In each resample, we refitted the control reference and primary four-level regional stage models, recalculated the standardised stage-coefficient map and scored it using the fixed contrast weights. We calculated the 95% percentile confidence interval for D from the 2.5th and 97.5th percentiles of the bootstrap distribution.

### Longitudinal mouse MRI after α-synuclein fibril injection

We obtained longitudinal regional structural MRI data for wild-type and hemizygous M83 mice injected in the right striatum with human or mouse α-synuclein preformed fibrils (PFFs), or phosphate-buffered saline (PBS)^49^. At each follow-up (30, 90 or 120 days after injection), we included mice with both a pre-injection baseline measurement and a measurement at that visit. Cohorts with only two imaging time points were excluded, following the source deformation-based morphometry analysis code.

The source measurements were regional means of unsmoothed Jacobian determinants that included linear and nonlinear transformations (dimensionless). We averaged corresponding left- and right-hemisphere measurements to obtain 211 bilateral region values. Each source region was assigned to the first structure represented in OTTER encountered when traversing the Allen ontology from that region towards its ancestors. Measurements from source regions assigned to the same OTTER structure were averaged with equal weight, and unmapped regions were excluded, yielding 205 structures.

At each follow-up and separately within genotype, we modelled each region’s MRI measurement as a function of its baseline value, sex and indicators for human and mouse PFF injection, with PBS as the reference group. Each PFF coefficient was negated and divided by the model residual standard deviation, so positive values indicated lower measurements in PFF-injected mice than in PBS controls. We z-standardised each resulting effect map across the 205 structures.

For each map, the interoceptive-minus-motor score was the mean across VISC, GU, AId, AIv and AIp minus the mean across primary and secondary motor cortex (MOp and MOs), with structures weighted equally within each set. At each follow-up, we averaged the scores for the four genotype-by-PFF conditions equally: wild-type/human PFF, wild-type/mouse PFF, M83/human PFF and M83/mouse PFF. We estimated the linear slope of this mean trajectory by ordinary least squares with an intercept, giving equal weight to the three follow-ups at 30, 90 and 120 days after injection.

We tested for a positive slope of the equally weighted mean trajectory using a one-sided animal-label permutation test, with 10,000 permutations for the primary analysis and normalisation sensitivities, and 5,000 for each model or sample sensitivity. Injection-group labels were permuted among animals within groups defined by genotype, sex and cohort. The batch-adjusted sensitivity additionally restricted permutations within acquisition batch. One reassigned label was retained across all scans from an animal, and both PFF comparisons used the shared PBS reference. We estimated 95% confidence intervals for the slope using 5,000 bootstrap resamples of animals, stratified by genotype, injection group and sex, retaining each sampled animal’s available visits together. After each permutation or bootstrap resample, we refitted the regional models and repeated residual-standard-deviation scaling, spatial normalisation, target scoring and calculation of the equally weighted mean slope. Confidence intervals were the 2.5th and 97.5th percentiles of the bootstrap slope estimates.

Sensitivity analyses added cohort indicators, added both cohort and acquisition-batch indicators, or combined cohort indicators with restriction to animals with all three follow-up scans. In separate normalisation sensitivities, we replaced whole-map z-standardisation with percentile ranks across all structures, or with z-scores or ranks calculated within isocortex. Ranks used average ranks for ties and were scaled as (rank − 0.5)/n, where n was the number of finite structures in the relevant set. Isocortical z-scores used the mean and sample standard deviation across isocortical structures. Slopes were expressed per day in the score units of each normalisation.

### Interface display categories

The released interface assigned each mouse ROI to one of five categories: anchor plus benchmark region, benchmark region only, anchor only, internally stable and other. Here, ‘anchor’ denoted membership in a point-anchor or regional correspondence set, and ‘benchmark region’ denoted membership in one of the 19 benchmark regions. The internal stability summary combined bootstrap stability of the highest-weight human partner, concentration of the ROI’s coupling weights and functional-connectivity similarity to a Garin point anchor.

These categories determined the spatial granularity displayed by the interface and were not used in inferential analyses. Category assignment did not provide calibrated confidence scores or estimates of individual ROI correspondence accuracy. In particular, membership in a benchmark region did not establish that the ROI’s own correspondence had been independently validated.

### Statistics and reproducibility

Correlations were assessed using Pearson’s r unless Spearman’s ρ was specified. Coupling-row and source-region-value permutation tests used 1,000 randomisations. Monte Carlo P values included a +1 correction. We controlled the false discovery rate at 5% using the Benjamini–Hochberg procedure, separately across the 19 benchmark pairs for transported-mass enrichment and for centroid displacement. The 16 network-versus-rest reconstruction contrasts used Bonferroni correction (adjusted P < 0.05). Figure 5c displays unadjusted P values for the seven reference-map comparisons; Results report which survived Benjamini–Hochberg correction across these seven tests at a 5% false discovery rate.

No statistical method was used to predetermine sample size for these analyses. Randomised analyses used fixed seeds specified in the released code.

## Code availability

The OTTER package, including data preparation, the model, and all analyses, is available at https://github.com/peach-lucien/otter under the MIT licence. It requires Python ≥ 3.10 and builds on the POT optimal-transport library (≥ 0.9.4)^64^ together with the standard scientific-Python and scverse packages.

## Data availability

Processed inputs, the canonical coupling, the outputs needed to reproduce the analyses and the full raw inputs for a from-scratch rebuild are archived on Zenodo (concept DOI 10.5281/zenodo.20733162, which resolves to the latest version; this study used version 1.3.0, DOI 10.5281/zenodo.21458106). The primary data derive from previously published resources cited above, including the Allen Mouse Connectivity Atlas, the Domhof et al. human connectome, the Allen Mouse Brain and Allen Human Brain atlases, the Garin homology atlas, the Beauchamp benchmark, and the TransBrain package with the Brainnetome atlas; their original terms of use apply. Restricted QPN-NC imaging and clinical data are available by application through Zenodo (10.5281/zenodo.17246063) where access is granted by the data owners on a case-by-case basis. The public longitudinal PFF source data are available from Zenodo (10.5281/zenodo.16614414).

## Acknowledgements

The authors would like to thank Clément M Garin and Suliann Ben Hamed from Institut des Sciences Cognitives Marc Jeannerod (UMR5229 CNRS Université de Lyon, Bron, France) for generous support and discussions related to the cross-species atlases of Garin et al.^23^.

This study was supported by the German Research Foundation (DFG, Project-ID 424778381-TRR 295 ReTune). Funding to SPK and PBS was provided by the German Federal Ministry of Research, Technology and Space (BMFTR, 01EJ2502A TAhRget and ERA-NET NEURON 01EW2305 IMatrix), and the DFG (EXC-2049-390688087 NeuroCure). C.W.I. receives funding from the Interdisciplinary Center for Clinical Research (IZKF) at the University of Würzburg (S-506, N-362) and by the VERUM foundation.

## Extended Data

**Extended Data Fig. 1.**
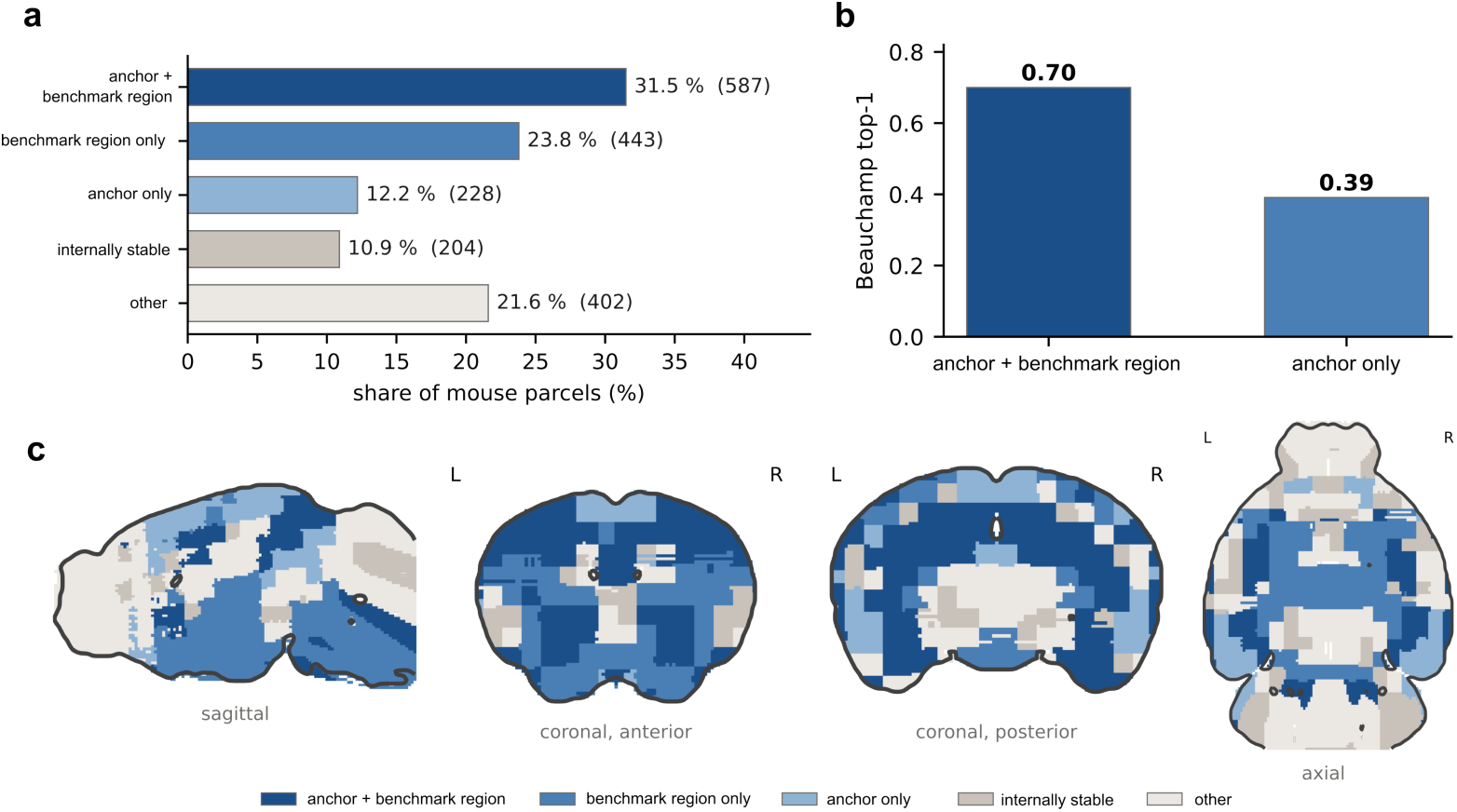
Mouse parcel categories based on anatomical supervision, benchmark coverage and internal stability. Mouse parcels are grouped into five categories using anatomical-anchor membership, inclusion in one of the 19 Beauchamp benchmark regions and internal stability (Methods). **a**, Percentage and number of the 1,864 mouse parcels in each category. **b**, Parcel-weighted regional top-1 recovery on the Beauchamp benchmark for the two categories containing benchmark-region parcels. **c**, Distribution of the categories across sagittal, coronal and axial mouse-brain sections.

**Extended Data Fig. 2.**
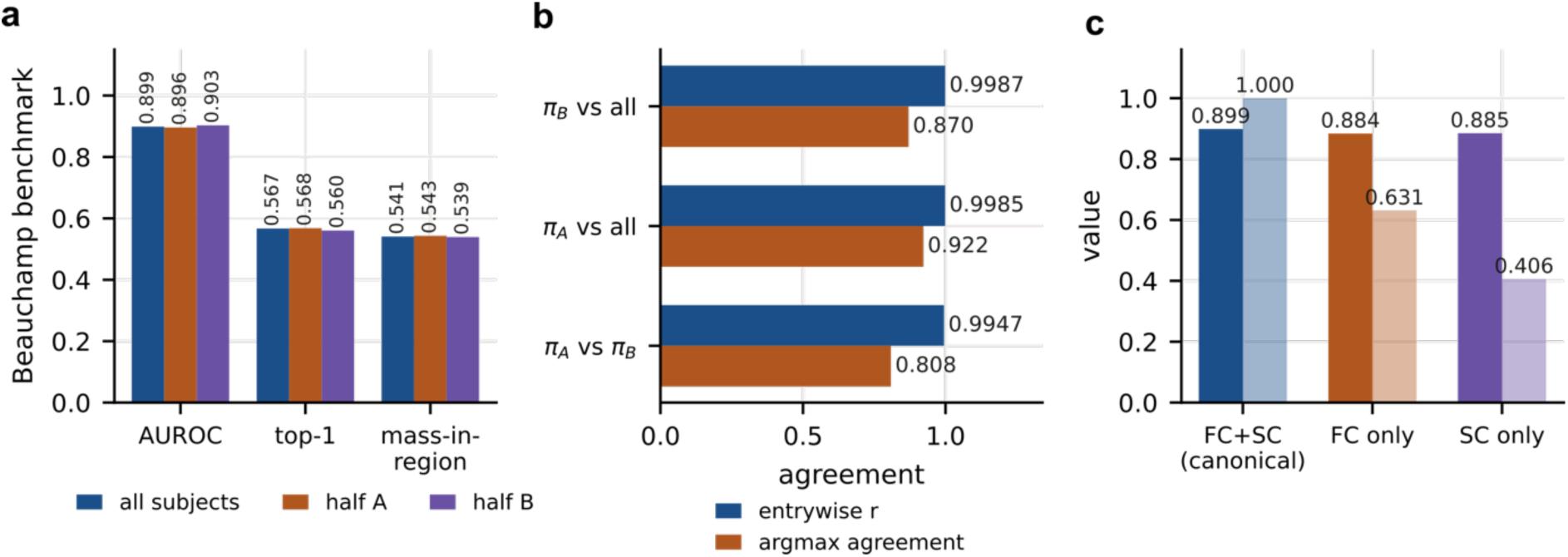
Effects of cohort splitting and connectivity modality on OTTER correspondence. **a**, Human (n = 113 participants) and mouse (n = 105 mice) functional-connectivity cohorts were randomly split into halves, and OTTER was refitted using the corresponding halves. Bars show regional AUROC, top-1 recovery and mass-in-region on the Beauchamp benchmark for the full cohorts (blue), half A (orange) and half B (purple). **b**, Pairwise agreement among the split-half couplings (πA and πB) and the full-cohort coupling. Blue bars show entrywise Pearson correlation; orange bars show the fraction of the 1,864 mouse parcels assigned to the same highest-weighted human target. **c**, Refits using both functional connectivity (FC) and structural connectivity (SC), FC alone or SC alone. Solid bars show regional AUROC; faded bars show the fraction of mouse parcels retaining the highest-weighted human target from the FC + SC model.

**Extended Data Fig. 3.**
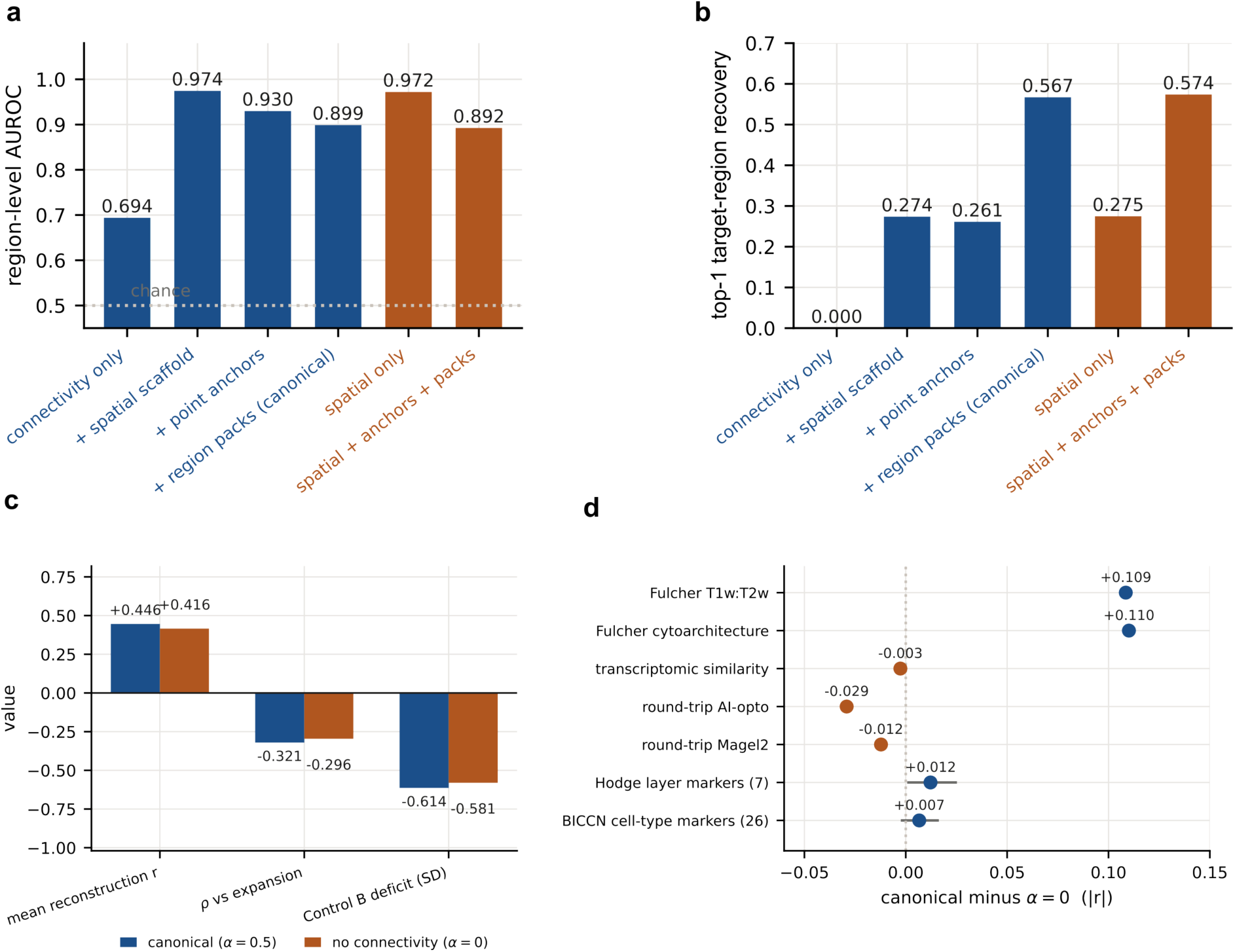
Effects of removing the connectivity cost from OTTER. **a**, Regional AUROC on the 19-region Beauchamp benchmark. Blue bars show connectivity alone, followed by successive addition of the spatial cost, point anchors and regional correspondence packs. Orange bars show fits without the connectivity cost (α = 0), using spatial information alone or combined with anatomical supervision. The spatial transformation was fitted to Garin landmarks. The dotted line indicates chance AUROC (0.5). **b**, Top-1 target-region recovery for the same fits. No target-wise withholding of anatomical supervision was applied in a or b. **c**, Comparison of the full model (blue, α = 0.5) and the model without connectivity (orange, α = 0): mean functional-connectivity reconstruction accuracy, the correlation between regional reconstruction accuracy and macaque-to-human cortical expansion (Spearman ρ), and the Control B reconstruction deficit (SD; *Fig. 5*). **d**, Differences in correlation magnitude for translated maps, transcriptomic similarity and round-trip consistency (full model minus α = 0). Blue and orange indicate positive and negative differences, respectively. For the two marker groups, points show mean differences and horizontal lines show 95% bootstrap confidence intervals.

**Extended Data Fig. 4.**
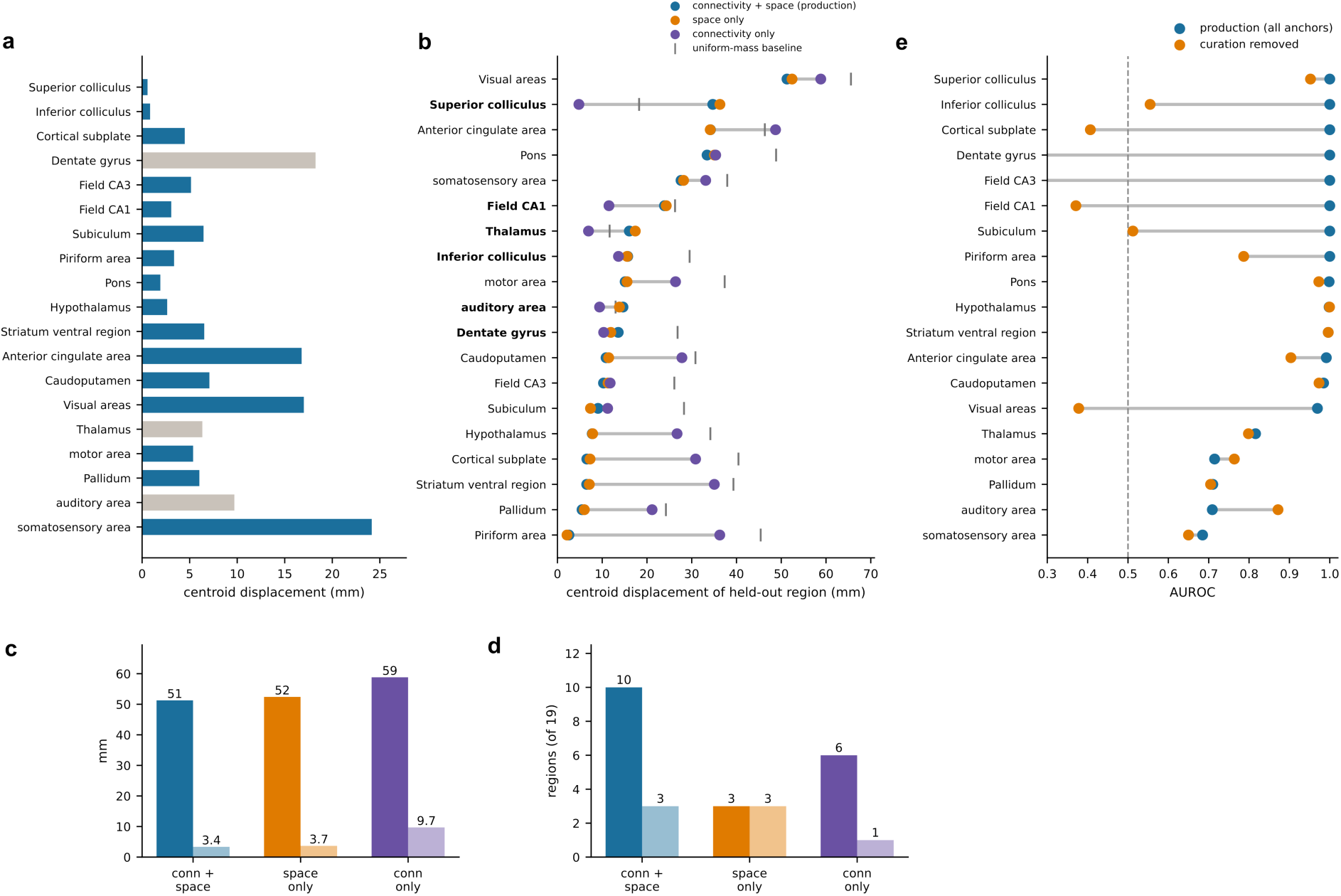
Regional localisation and recovery on the Beauchamp benchmark. **a**, Full-model localisation error for the 19 region pairs, measured as the distance between the mass-weighted centroid of the predicted human distribution and the reference human-region centroid (mm). Blue bars indicate significance against a target-centroid rotation null (FDR q < 0.05). **b**, Localisation error after withholding each region’s overlapping anatomical supervision and refitting with connectivity and spatial costs (blue), spatial cost without connectivity (orange), or connectivity without the spatial cost (purple). Supervision elsewhere was retained. Grey ticks show region-specific baseline errors for uniformly distributed mass over human parcels (mean, 33.2 mm; range, 11.7–65.6 mm). Bold labels identify regions where omitting the spatial cost gave the lowest error. **c**, Maximum regional error (solid bars) and mean additional error relative to each region’s best-performing configuration, averaged across the 19 regions (faded bars). **d**, Number of regions with the lowest error (solid bars) or error greater than their uniform-mass baseline (faded bars). Colours in c and d match b. **e**, Regional AUROC for the full model (blue) and refits withholding each region’s overlapping supervision (orange). Parcel-weighted AUROC decreased from 0.90 to 0.73, with five of 19 held-out regions below chance (dashed line, 0.5). Grey lines in b and e connect estimates for the same region.

**Extended Data Fig. 5.**
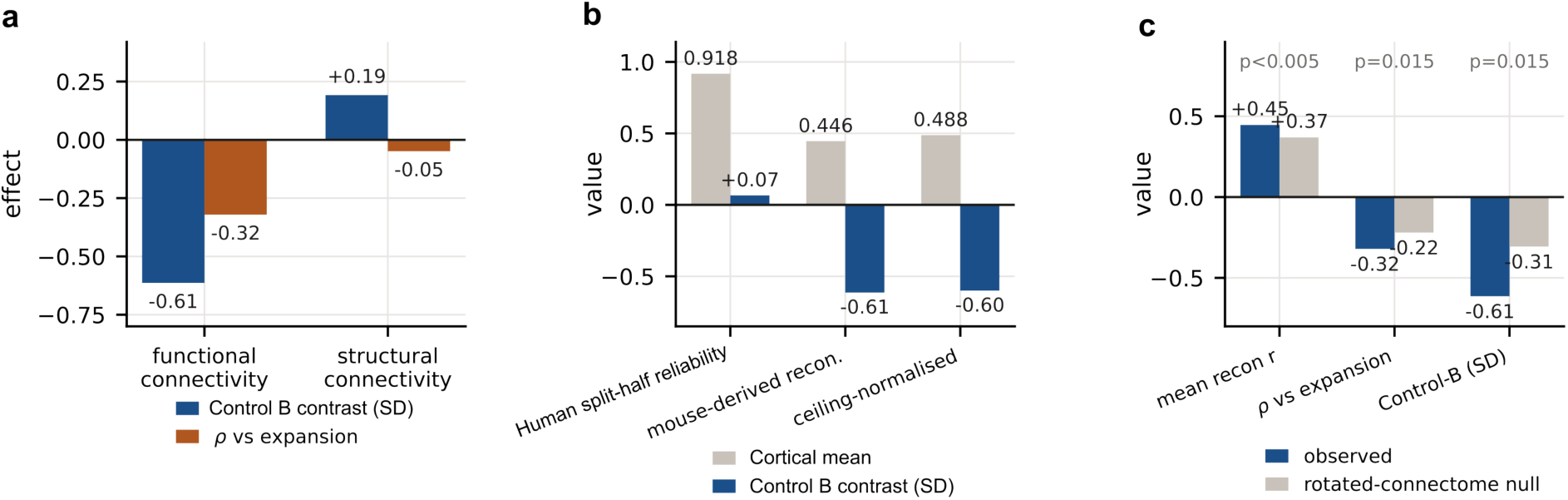
Controls for regional differences in mouse-based connectivity reconstruction. All analyses use the same fitted OTTER coupling. Control B contrasts compare the mean within this subdivision of the frontoparietal control network with the mean across the remaining cortex, expressed in units of each map’s cortical standard deviation. **a**, Control B contrast (blue) and Spearman correlation between reconstruction accuracy and macaque-to-human cortical expansion^14^ (orange), calculated using either functional or structural connectomes from both species. The Control B contrast for functional reconstruction is reported descriptively because it did not survive correction across the 16 networks tested. **b**, Human split-half reliability, mouse-based reconstruction accuracy and reconstruction accuracy divided by split-half reliability at each human parcel. Reliability was calculated by correlating each parcel’s connectivity profiles estimated from two halves of the human cohort. Grey bars show cortical means; blue bars show Control B contrasts. **c**, Observed values (blue) and means across 200 spatial surrogate mouse connectomes (grey), compared for mean reconstruction accuracy, expansion correlation and Control B contrast. Surrogates were generated by rotating mouse parcel coordinates and reassigning connectome rows and columns according to the nearest original parcel coordinates. P-values compare the observed statistics with their corresponding surrogate distributions.

**Extended Data Fig. 6.**
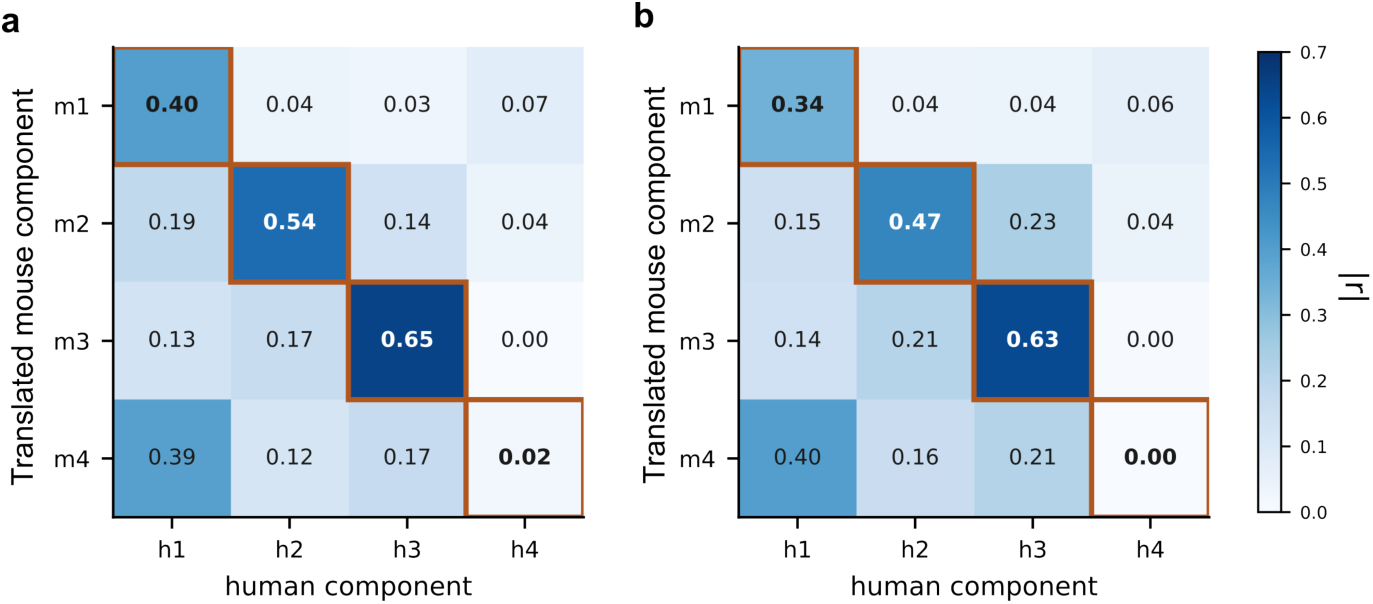
Transfer of functional connectivity gradients between mouse and human. The first four diffusion-map components were derived separately from the mouse and human functional connectomes. Each mouse component (rows, m1–m4) was translated to human parcels and correlated with each human component (columns, h1–h4). **a**, Full OTTER coupling. **b**, Coupling refitted without the connectivity cost (α = 0). Numbers and colours show absolute correlations (|r|); orange outlines mark pairs with the same component index. In both panels, mouse components 1–3 had their highest absolute correlations with the same-index human components, whereas mouse component 4 had its highest absolute correlation with human component 1.

**Extended Data Fig. 7.**
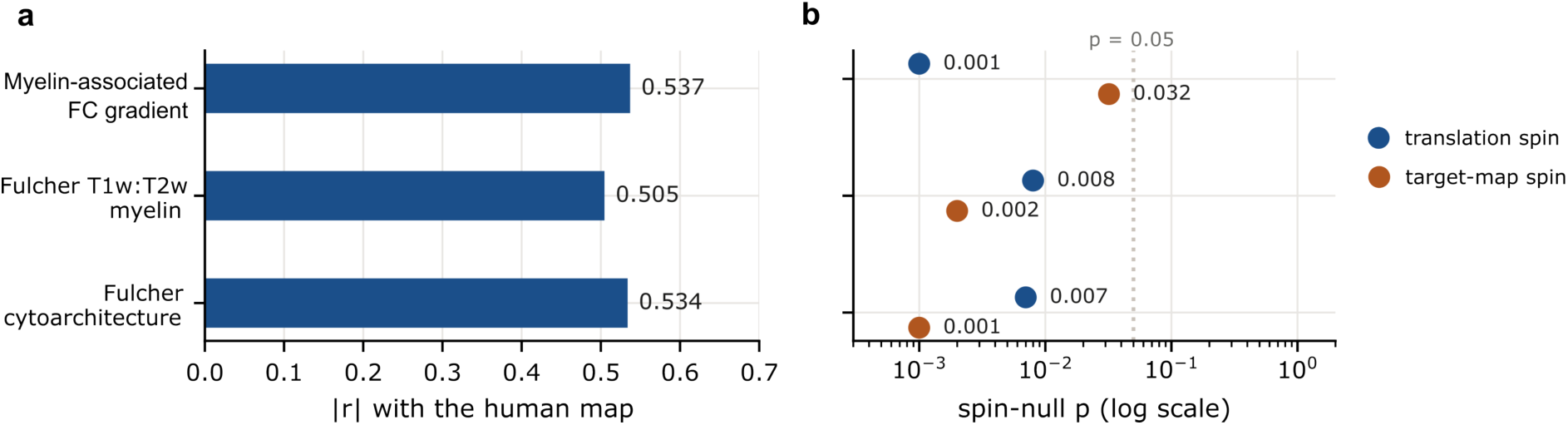
Spatial-rotation tests of mouse-to-human map correspondence. **a**, Absolute Pearson correlations (|r|) between translated mouse maps and their human comparison maps. The myelin-associated functional-connectivity gradient was compared with the corresponding human gradient; the mouse T1w/T2w ratio and cytoarchitectural-type maps were both compared with the human T1w/T2w ratio. Comparisons included 2,094, 1,789 and 1,787 human parcels, respectively; these counts describe the size of each spatial comparison rather than independent observations. **b**, Unadjusted empirical P-values on a logarithmic scale for the translation-spin (blue) and target-map spin (orange) tests. The translation-spin test rotates the mouse map before translation through the fixed OTTER coupling. The target-map spin test rotates the human comparison map while keeping the translated mouse map fixed. Both tests used 1,000 uniformly random whole-sphere rotations with nearest-neighbour parcel reassignment. P-values were calculated as (B + 1)/1,001, where B is the number of rotations with an absolute correlation at least as large as observed. The dotted line marks P = 0.05.

**Extended Data Fig. 8.**
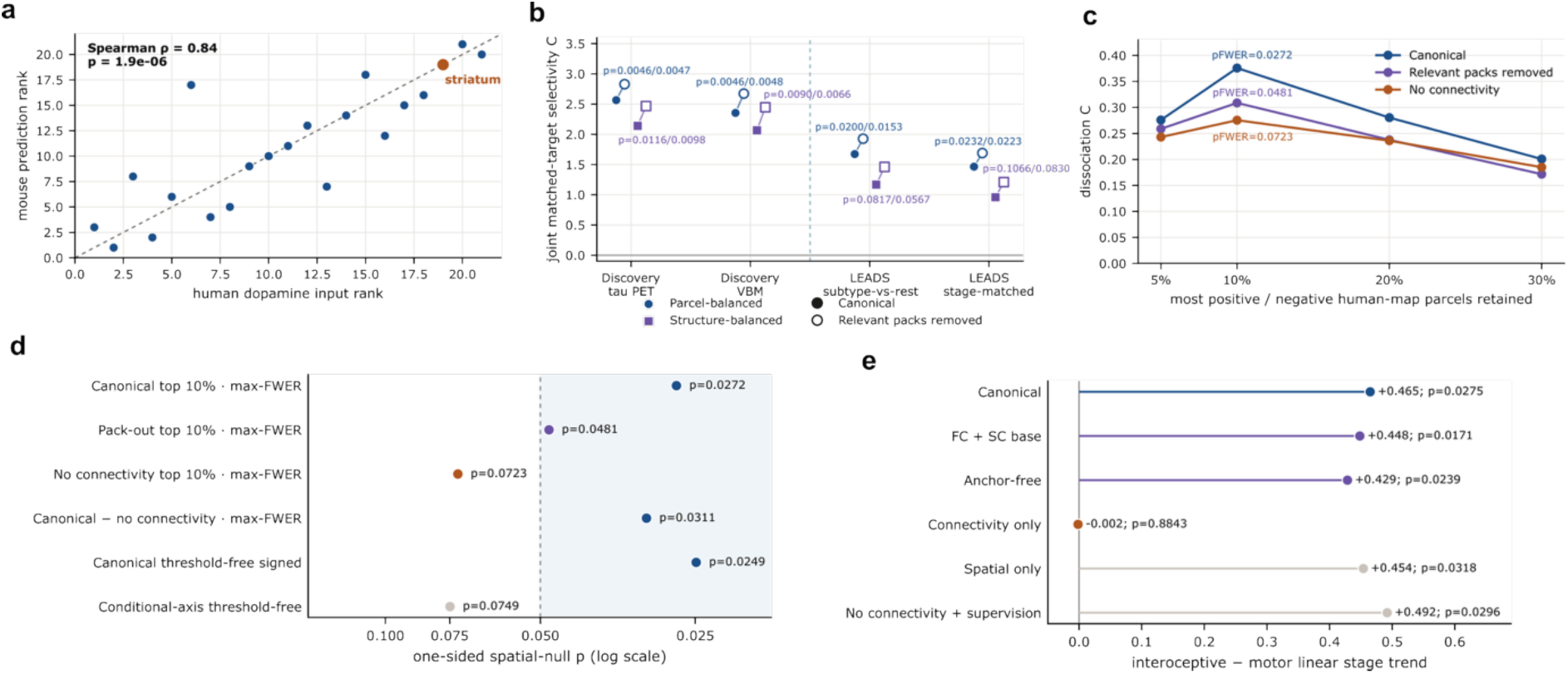
Dopamine map correspondence and sensitivity analyses of human-to-mouse translation. **a**, Ranks of mean human FP-CIT SPECT input values and mean translated mouse values across 21 paired anatomical classes, defined by each parcel’s nearest same-hemisphere point anchor (Spearman ρ = 0.84; two-sided asymptotic P = 1.9 × 10⁻⁶). Orange, striatum; dashed line, equal ranks. **b**, Alzheimer phenotype joint selectivity, calculated by summing each phenotype’s assigned-target score minus the mean of its two alternative-target scores. Blue circles weight mouse parcels equally within each target set; purple squares weight anatomical structures equally. Filled points show full OTTER; open points show refits without the relevant regional correspondence packs. Lines connect full-model and refitted estimates for the same dataset and weighting. Labels give full-model/refit P-values (one-sided; 10,000 spatial rotations). **c**, Transcranial magnetic stimulation (TMS) circuit separation across the 5%, 10%, 20% and 30% positive and negative human-map tails. For each translated map, bias is the medial prefrontal/cingulate score minus the amygdala/insula score; C is dysphoric bias minus anxiosomatic bias. Blue, full OTTER; purple, relevant packs removed; orange, matched coupling without the connectivity cost. Labels at the primary 10% threshold give one-sided P values with maximum-statistic family-wise-error correction across the four thresholds. **d**, One-sided spatial-null P values for the listed TMS comparisons, including the paired difference between full OTTER and the coupling without connectivity. The first four rows are each corrected across the four thresholds; the last two are threshold-free analyses. The primary TMS null used 10,000 Moran spectral randomisations. Dashed line, P = 0.05; shading, P < 0.05 (logarithmic scale). **e**, Linear stage trends in translated scores for visceral, gustatory and agranular-insular regions relative to primary motor cortex, estimated from cross-sectional Parkinson cortical-thickness maps under alternative couplings. Labels give two-sided spatial-null P-values. Positive trends were also obtained without connectivity, whereas connectivity alone gave an estimate near zero.

**Extended Data Fig. 9.**
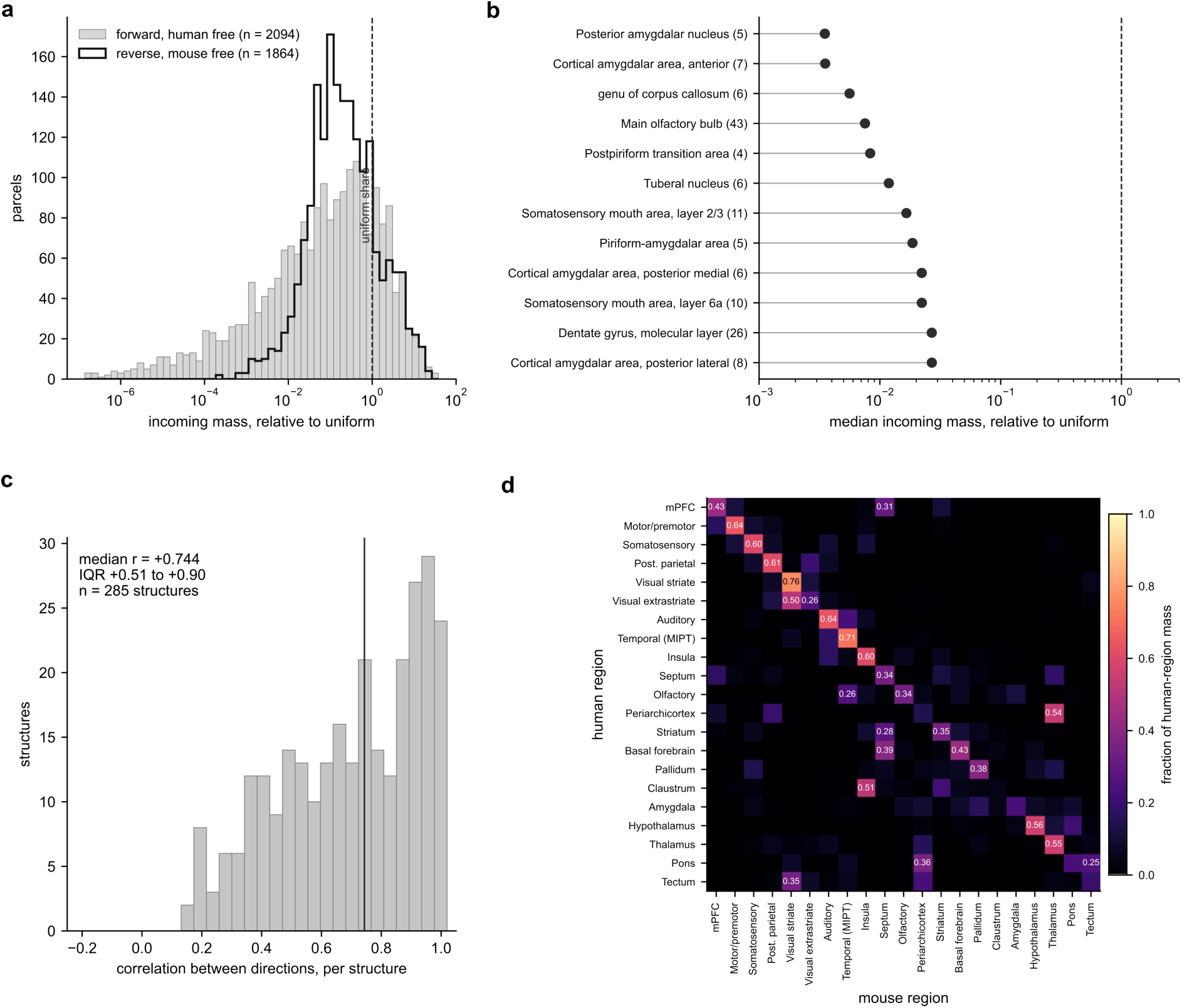
Refitting OTTER with a fixed human marginal distribution. The original fit constrains all mouse parcels to have the same total correspondence weight, while human parcel totals are unconstrained. Here, the model was refitted with human parcel totals fixed uniformly and mouse parcel totals unconstrained, using the same connectivity and cross-species costs. **a**, Received correspondence weight per parcel, divided by the uniform share for the unconstrained species. Grey, human parcels in the original fit (n = 2,094); black outline, mouse parcels in the human-fixed refit (n = 1,864). The dashed line marks a relative weight of 1, corresponding to uniform allocation. The proportions receiving less than 0.1 were 50.4% and 35.9%, respectively. **b**, The twelve mouse structures with the lowest median received weight in the human-fixed refit, among 148 structures represented by at least four parcels. Weights are normalised as in a; numbers in parentheses give parcel counts for that structure. The dashed line marks the uniform reference. Horizontal axes in a and b are logarithmic. **c**, Correlations between each mouse structure’s distribution of correspondence weight across human parcels under the two fits (285 structures). The vertical line marks the median (r = 0.744; interquartile range, 0.51–0.90). **d**, The human-fixed coupling aggregated to 21 cross-species homology classes and normalised within each human class. Each row shows the fraction of that human class’s weight assigned to each mouse class. The mean fraction assigned to the corresponding class was 0.43; equal allocation across the 21 mouse classes would give 0.048.

**Extended Data Fig. 10.**
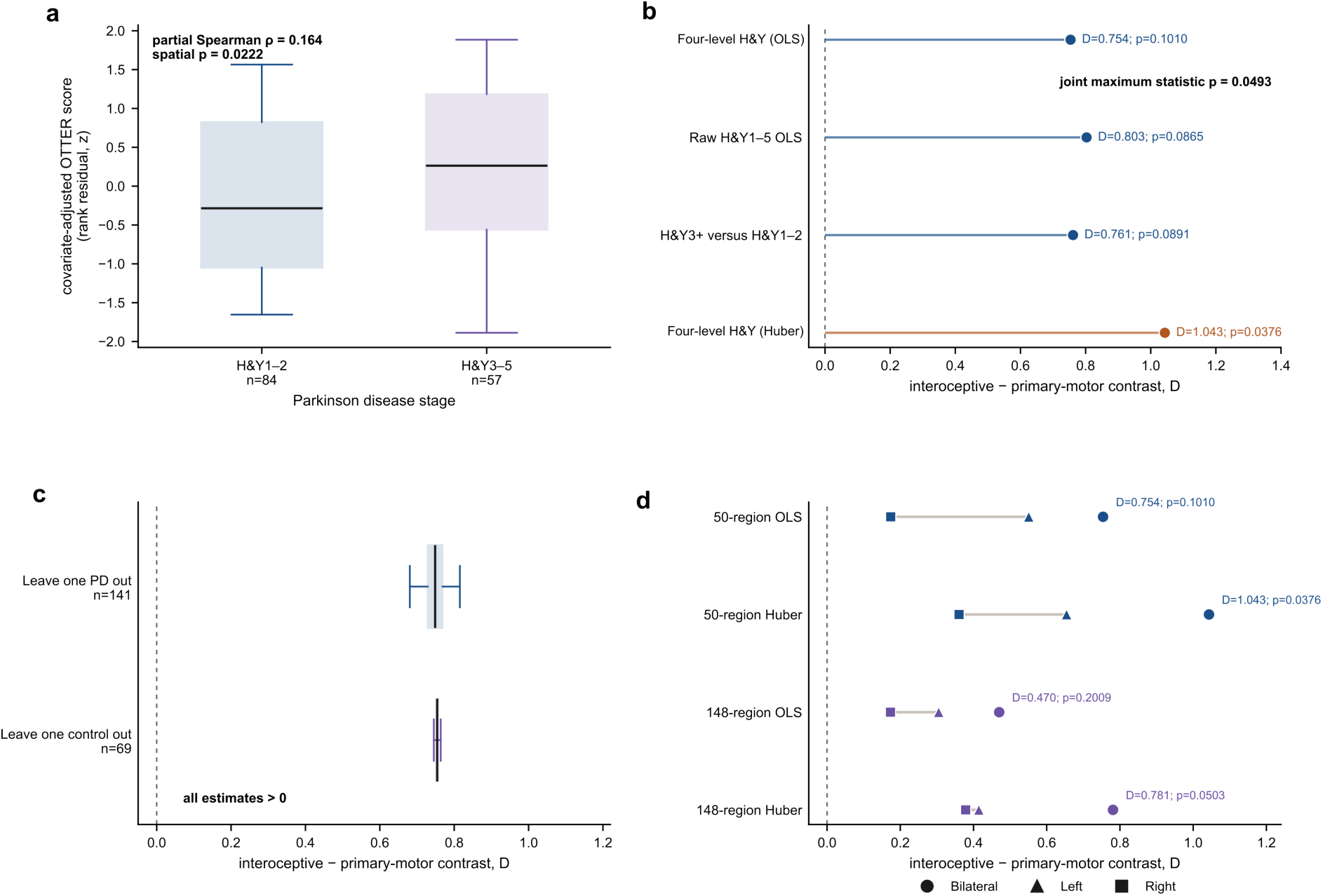
Parkinson disease stage and translated cortical thinning in an independent cohort. **a**, Cortical-thinning maps from 141 participants with Parkinson disease in the Quebec Parkinson Network Neuroimaging Cohort were estimated relative to 69 controls and translated into scores comparing visceral, gustatory and agranular-insular regions (the interoceptive set) with primary motor cortex. Boxplots show standardised, covariate-adjusted rank residuals for Hoehn–Yahr (H&Y) stages 1–2 (n = 84) and 3–5 (n = 57). The partial Spearman correlation uses all four ordinal levels (1, 2, 3 and 4/5), adjusting for MRI age, sex and clinical–MRI assessment lag (ρ = 0.164, P = 0.0222). **b**, D is the mean translated stage-association score in the interoceptive set minus that in primary motor cortex, calculated after standardising the human regional coefficient map. The three ordinary least-squares (OLS) models use four-level stage, raw stages 1–5, or stages 3–5 versus 1–2; Huber regression uses the four-level stage coding. A maximum-statistic test across these four regional models and the participant-level analysis gave P = 0.0493. **c**, Distributions of D after excluding each participant with Parkinson disease (141 refits) or each control (69 refits). All estimates remained positive. **d**, D estimated using 50 cortical regions shared by the cohort and OTTER (blue) or 148 native Destrieux regions (purple), under OLS and Huber regression. Circles, bilateral; triangles, left hemisphere; squares, right hemisphere. Lines connect hemisphere-specific estimates. All estimates were positive. Boxplots in a and c show medians, interquartile ranges and whiskers extending to the most extreme values within 1.5 times the interquartile range; outlier markers are omitted. Spatial P-values use one-sided tests with 10,000 hemisphere-preserving, one-to-one rotations.

**Extended Data Fig. 11.**
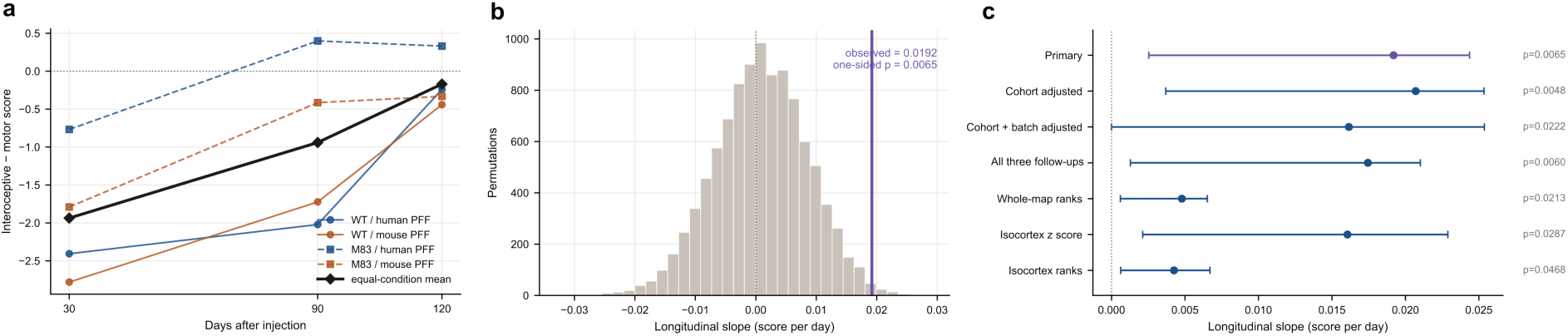
Longitudinal comparison of atrophy in interoceptive and motor regions after α-synuclein fibril injection. Wild-type and hemizygous M83 mice received right-striatal human PFF, mouse PFF or PBS injections and underwent baseline and follow-up structural MRI. Totals across genotype and treatment groups were 217, 215 and 88 animals at 30, 90 and 120 days, respectively. **a**, The score is the mean standardised atrophy across visceral, gustatory and the three agranular-insular subdivisions minus the mean across primary and secondary motor cortex, weighting structures equally. Regional PFF effects were estimated relative to genotype-matched PBS controls, adjusted for baseline volume and sex, scaled by the residual standard deviation and z-standardised across 205 structures. Coloured lines show the four genotype-by-PFF conditions; black diamonds show their equally weighted mean. **b**, Slope null distribution from 10,000 animal-level treatment-label permutations within genotype, sex and cohort, retaining each animal’s label across visits. The purple line marks the observed slope (0.0192 score units per day; P = 0.0065). **c**, Slopes of the equally weighted condition mean and 95% confidence intervals from 5,000 animal-bootstrap resamples stratified by genotype, injection and sex. Sensitivities adjust for cohort and acquisition batch, restrict the sample to animals with all three follow-ups, or alter effect-map normalisation. Slopes are expressed per day using each analysis’s score units. P-values are one-sided animal-label permutation tests, using 10,000 permutations for the primary analysis and 5,000 for each sensitivity. PFF, preformed fibril; PBS, phosphate-buffered saline; M83, hemizygous A53T α-synuclein transgenic mice; WT, wild type.

---

**Supplementary Table 1.**
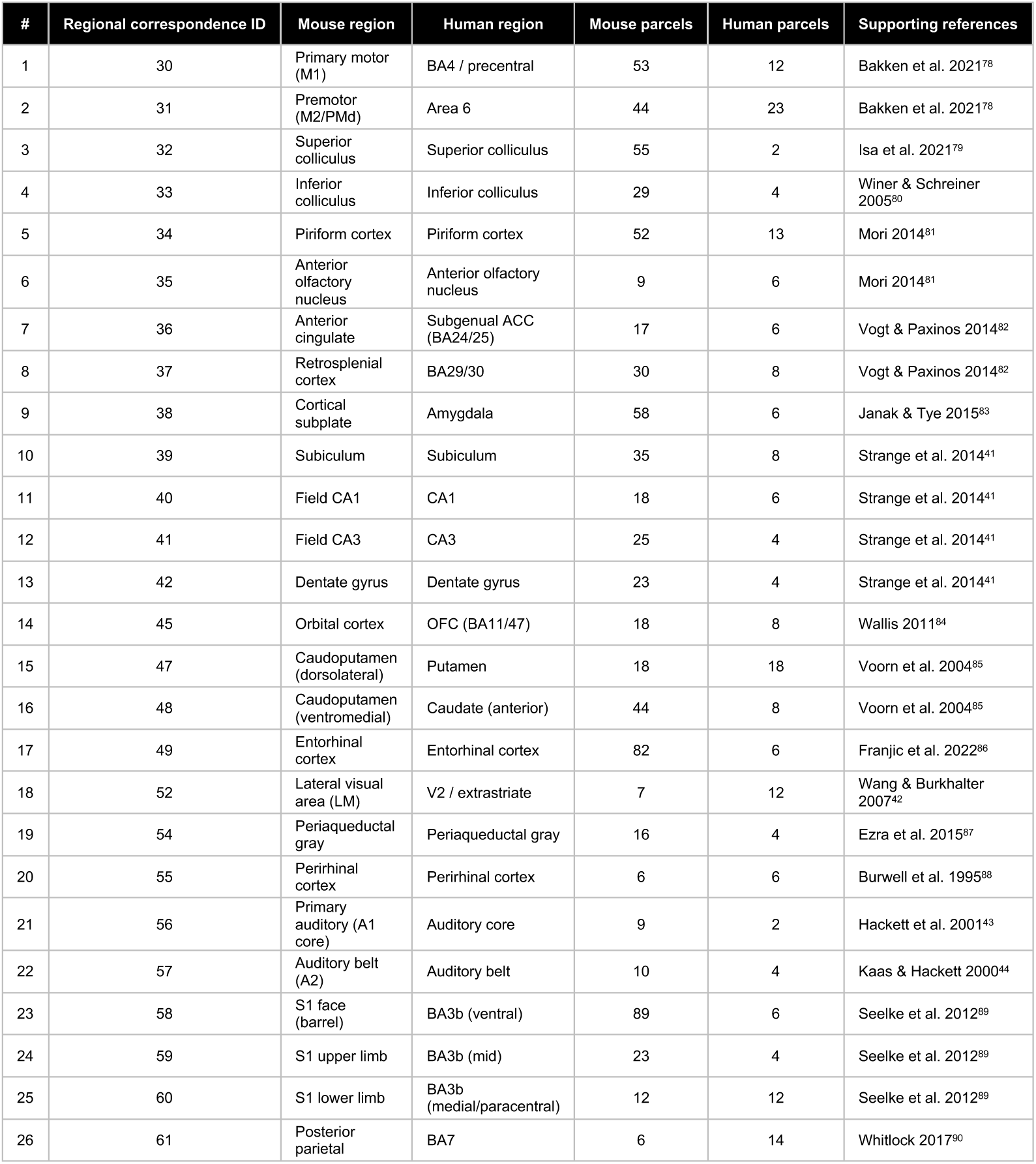
Regional anatomical correspondences used to fit OTTER. The table lists 26 mouse–human regional correspondences curated from the cited literature and used as soft anatomical constraints when fitting OTTER. These regional constraints supplement the point landmarks derived from the Garin atlas^23^. Each correspondence includes parcels from both hemispheres; counts refer to the 1,864-parcel mouse and 2,094-parcel human parcellations used for fitting. BA (Brodmann area), ACC (anterior cingulate cortex), OFC (orbitofrontal cortex) and S1 (primary somatosensory cortex).

| # | Regional correspondence ID | Mouse region | Human region | Mouse parcels | Human parcels | Supporting references |
|---|---|---|---|---|---|---|
| 1 | 30 | Primary motor (M1) | BA4 / precentral | 53 | 12 | Bakken et al. 2021<sup>78</sup> |
| 2 | 31 | Premotor (M2/PMd) | Area 6 | 44 | 23 | Bakken et al. 2021<sup>78</sup> |
| 3 | 32 | Superior colliculus | Superior colliculus | 55 | 2 | Isa et al. 2021<sup>79</sup> |
| 4 | 33 | Inferior colliculus | Inferior colliculus | 29 | 4 | Winer & Schreiner 2005<sup>80</sup> |
| 5 | 34 | Piriform cortex | Piriform cortex | 52 | 13 | Mori 2014<sup>81</sup> |
| 6 | 35 | Anterior olfactory nucleus | Anterior olfactory nucleus | 9 | 6 | Mori 2014<sup>81</sup> |
| 7 | 36 | Anterior cingulate | Subgenual ACC (BA24/25) | 17 | 6 | Vogt & Paxinos 2014<sup>82</sup> |
| 8 | 37 | Retrosplenial cortex | BA29/30 | 30 | 8 | Vogt & Paxinos 2014<sup>82</sup> |
| 9 | 38 | Cortical subplate | Amygdala | 58 | 6 | Janak & Tye 2015<sup>83</sup> |
| 10 | 39 | Subiculum | Subiculum | 35 | 8 | Strange et al. 2014<sup>41</sup> |
| 11 | 40 | Field CA1 | CA1 | 18 | 6 | Strange et al. 2014<sup>41</sup> |
| 12 | 41 | Field CA3 | CA3 | 25 | 4 | Strange et al. 2014<sup>41</sup> |
| 13 | 42 | Dentate gyrus | Dentate gyrus | 23 | 4 | Strange et al. 2014<sup>41</sup> |
| 14 | 45 | Orbital cortex | OFC (BA11/47) | 18 | 8 | Wallis 2011<sup>84</sup> |
| 15 | 47 | Caudoputamen (dorsolateral) | Putamen | 18 | 18 | Voorn et al. 2004<sup>85</sup> |
| 16 | 48 | Caudoputamen (ventromedial) | Caudate (anterior) | 44 | 8 | Voorn et al. 2004<sup>85</sup> |
| 17 | 49 | Entorhinal cortex | Entorhinal cortex | 82 | 6 | Franjic et al. 2022<sup>86</sup> |
| 18 | 52 | Lateral visual area (LM) | V2 / extrastriate | 7 | 12 | Wang & Burkhalter 2007<sup>42</sup> |
| 19 | 54 | Periaqueductal gray | Periaqueductal gray | 16 | 4 | Ezra et al. 2015<sup>87</sup> |
| 20 | 55 | Perirhinal cortex | Perirhinal cortex | 6 | 6 | Burwell et al. 1995<sup>88</sup> |
| 21 | 56 | Primary auditory (A1 core) | Auditory core | 9 | 2 | Hackett et al. 2001<sup>43</sup> |
| 22 | 57 | Auditory belt (A2) | Auditory belt | 10 | 4 | Kaas & Hackett 2000<sup>44</sup> |
| 23 | 58 | S1 face (barrel) | BA3b (ventral) | 89 | 6 | Seelke et al. 2012<sup>89</sup> |
| 24 | 59 | S1 upper limb | BA3b (mid) | 23 | 4 | Seelke et al. 2012<sup>89</sup> |
| 25 | 60 | S1 lower limb | BA3b (medial/paracentral) | 12 | 12 | Seelke et al. 2012<sup>89</sup> |
| 26 | 61 | Posterior parietal | BA7 | 6 | 14 | Whitlock 2017<sup>90</sup> |

**Supplementary Table 2.**
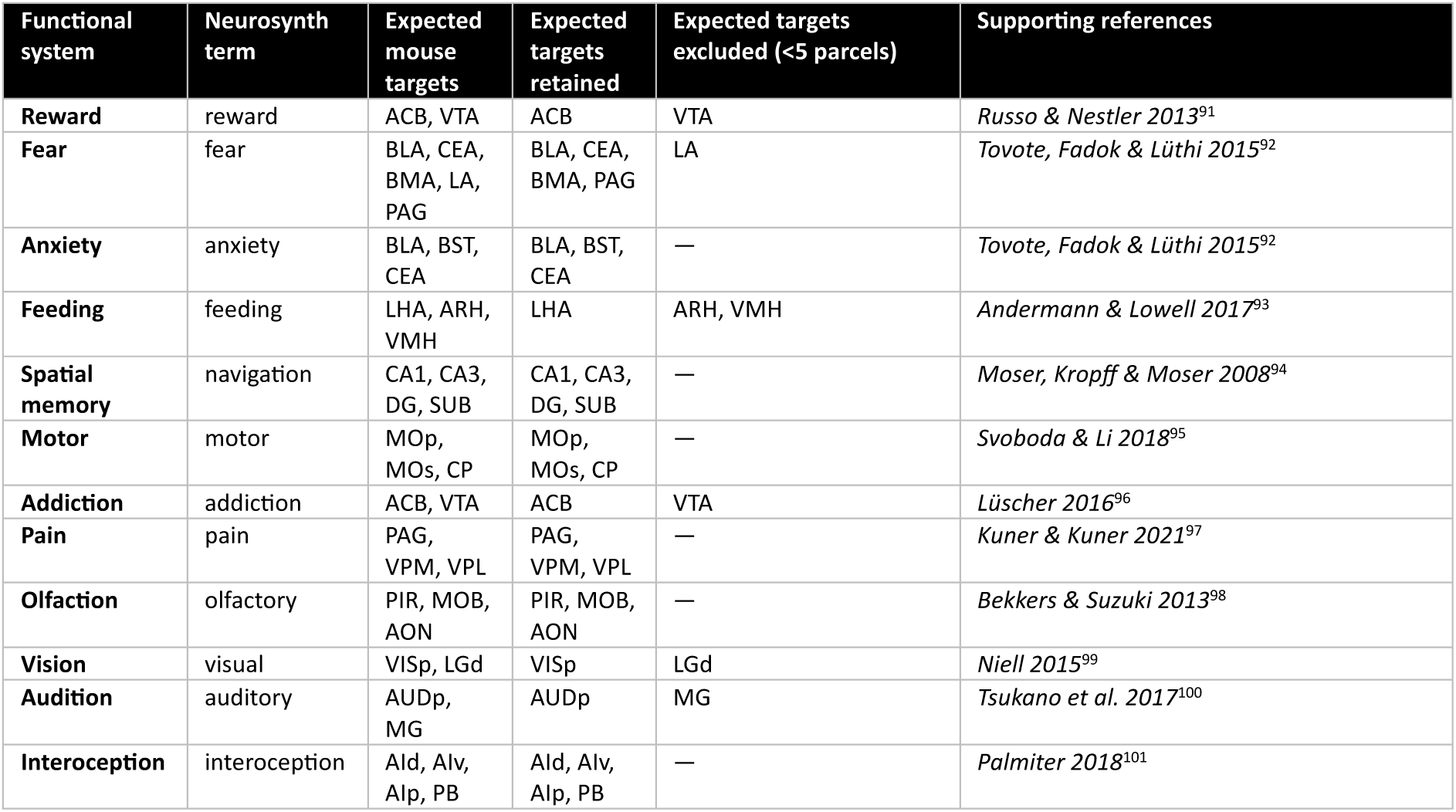
Expected mouse targets for evaluating human-to-mouse translation. The table lists the Neurosynth terms and expected mouse structures for the twelve functional systems evaluated in *Fig. 6d*. Expected targets were specified before scoring. Each translated map was used to rank 83 mouse structures represented by at least five parcels. Six distinct expected structures were excluded by this criterion, leaving only cortical targets for vision and audition.

| Functional system | Neurosynth term | Expected mouse targets | Expected targets retained | Expected targets excluded (<5 parcels) | Supporting references |
|---|---|---|---|---|---|
| **Reward** | reward | ACB, VTA | ACB | VTA | *Russo & Nestler 2013*<sup>91</sup> |
| **Fear** | fear | BLA, CEA, BMA, LA, PAG | BLA, CEA, BMA, PAG | LA | *Tovote, Fadok & Lüthi 2015*<sup>92</sup> |
| **Anxiety** | anxiety | BLA, BST, CEA | BLA, BST, CEA | — | *Tovote, Fadok & Lüthi 2015*<sup>92</sup> |
| **Feeding** | feeding | LHA, ARH, VMH | LHA | ARH, VMH | *Andermann & Lowell 2017*<sup>93</sup> |
| **Spatial memory** | navigation | CA1, CA3, DG, SUB | CA1, CA3, DG, SUB | — | *Moser, Kropff & Moser 2008*<sup>94</sup> |
| **Motor** | motor | MOp, MOs, CP | MOp, MOs, CP | — | *Svoboda & Li 2018*<sup>95</sup> |
| **Addiction** | addiction | ACB, VTA | ACB | VTA | *Lüscher 2016*<sup>96</sup> |
| **Pain** | pain | PAG, VPM, VPL | PAG, VPM, VPL | — | *Kuner & Kuner 2021*<sup>97</sup> |
| **Olfaction** | olfactory | PIR, MOB, AON | PIR, MOB, AON | — | *Bekkers & Suzuki 2013*<sup>98</sup> |
| **Vision** | visual | VISp, LGd | VISp | LGd | *Niell 2015*<sup>99</sup> |
| **Audition** | auditory | AUDp, MG | AUDp | MG | *Tsukano et al. 2017*<sup>100</sup> |
| **Interoception** | interoception | AId, AIv, AIp, PB | AId, AIv, AIp, PB | — | *Palmiter 2018*<sup>101</sup> |

